# Polar Crystal Habit and 3D Electron Diffraction Reveal the Malaria Pigment Hemozoin as a Selective Mixture of Centrosymmetric and Chiral Stereoisomers

**DOI:** 10.1101/2022.09.15.507960

**Authors:** Paul Benjamin Klar, David Waterman, Tim Gruene, Debakshi Mullick, Yun Song, James B. Gilchrist, C. David Owen, Wen Wen, Idan Biran, Lothar Houben, Neta Regev-Rudzki, Ron Dzikowski, Noa Marom, Lukas Palatinus, Peijun Zhang, Leslie Leiserowitz, Michael Elbaum

## Abstract

Detoxification of heme in Plasmodium depends on its crystallization into hemozoin. This pathway is a major target of antimalarial drugs. X-ray powder diffraction has established that the unit cell contains a cyclic hematin dimer, yet the pro-chiral nature of heme supports formation of four distinct stereoisomers, two centrosymmetric and two chiral enantiomers. Here we apply emerging methods of in situ cryo-electron tomography and diffraction to obtain a definitive structure of biogenic hemozoin. Individual crystals take a striking polar morphology. Diffraction analysis, supported by density functional theory, indicates a compositional mixture of one centrosymmetric and one chiral dimer, whose absolute configuration has been determined on the basis of crystal morphology and interaction with the aqueous medium. Structural modeling of the heme detoxification protein suggests a mechanism for dimer selection. The refined structure of hemozoin should serve as a guide to new drug development.

---

Malaria is a deadly disease that remains endemic to much of the world despite extraordinary efforts to its eradication ^[1]^. Anopheles mosquitos are the universal vector, and several species of Plasmodium affect humans, where they infect the red blood cells and catabolize the hemoglobin protein as a source of biomolecules. The heme so released is detoxified by sequestration into physiologically insoluble crystals of hemozoin, also known diagnostically as the malaria pigment. Disease symptoms include anemia and severe paroxysms resulting from immune response to hemozoin released to the blood stream ^[2]^. Growth of hemozoin is a target of common antimalarial drugs ^[3–6]^. Therefore, a precise determination of the crystal structure and morphology is of prime medical importance as well as fundamental interest.

A landmark study by X-ray powder diffraction (XRPD) revealed the crystalline structure of synthetic hemozoin, known as β-hematin ^[7]^. The unit cell contains not the monomeric heme present in hemoglobin, but rather a cyclic dimer. The structure was refined in a triclinic 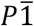 arrangement with one dimer per unit cell. Cyclic H-bonds between propionic acid groups form chains linking the hematin dimers along the ***a***−***c*** direction. The growth morphology is rod-like with well-developed {100} faces and {010} side faces of variable width, capped by slanted {011} end faces, as determined by transmission electron microscopy and diffraction^[8]^. Morphology of the synthetic crystal is in agreement with a theoretical growth form based on attachment energies ^[9]^.

While the model structure of β-hematin was centrosymmetric, the pro-chiral nature of the heme supports four configurations of the dimer ^[9–12]^. Specifically, methyl – vinyl pairs opposite the propionic acids are arranged asymmetrically, defining *re* (*R, rectus*) and *si* (*S, sinister*) heme faces. Heme binding proteins typically distinguish these two inequivalent orientations ^[13]^. Cyclic dimers form distinct isomers depending on which heme faces are brought to contact: *R*/*S*′, *S*/*R*′, *R*/*R*′, or *S*/*S*′. The first two are centrosymmetric, whereas the latter two are chiral enantiomers with pseudo-centrosymmetry (Fig. 1, Fig. S3). Straasø et al. interpreted XRPD data of synthetic hemozoin in terms of a major and minor centrosymmetric phase and suggested that these represent *R*/*S*′ and *S*/*R*′ isoforms, each containing additionally *R*/*R*′ and *S*/*S*′ dimers^[12]^ in significant concentration ^[11]^. Bohle et al. then reanalyzed the earlier powder diffraction and interpreted the data in terms of a disordered structure of *R*/*S*′ and *S*/*R*′ dimers in a 3:1 ratio ^[14]^. Complementary evidence that synthetic hemozoin is comprised of a mixture of the different isomeric hematin dimers also stemmed from a single-crystal X-ray diffraction study of a hematin dimer-DMSO solvate ^[15]^. More recently, a study of synthetic hemozoin by X-ray free electron laser diffraction ^[16]^ suggested that the structure of nano-crystals may differ from that of larger one due to greater conformational flexibility in the propionic chains ^[17]^.

**Figure 1.**
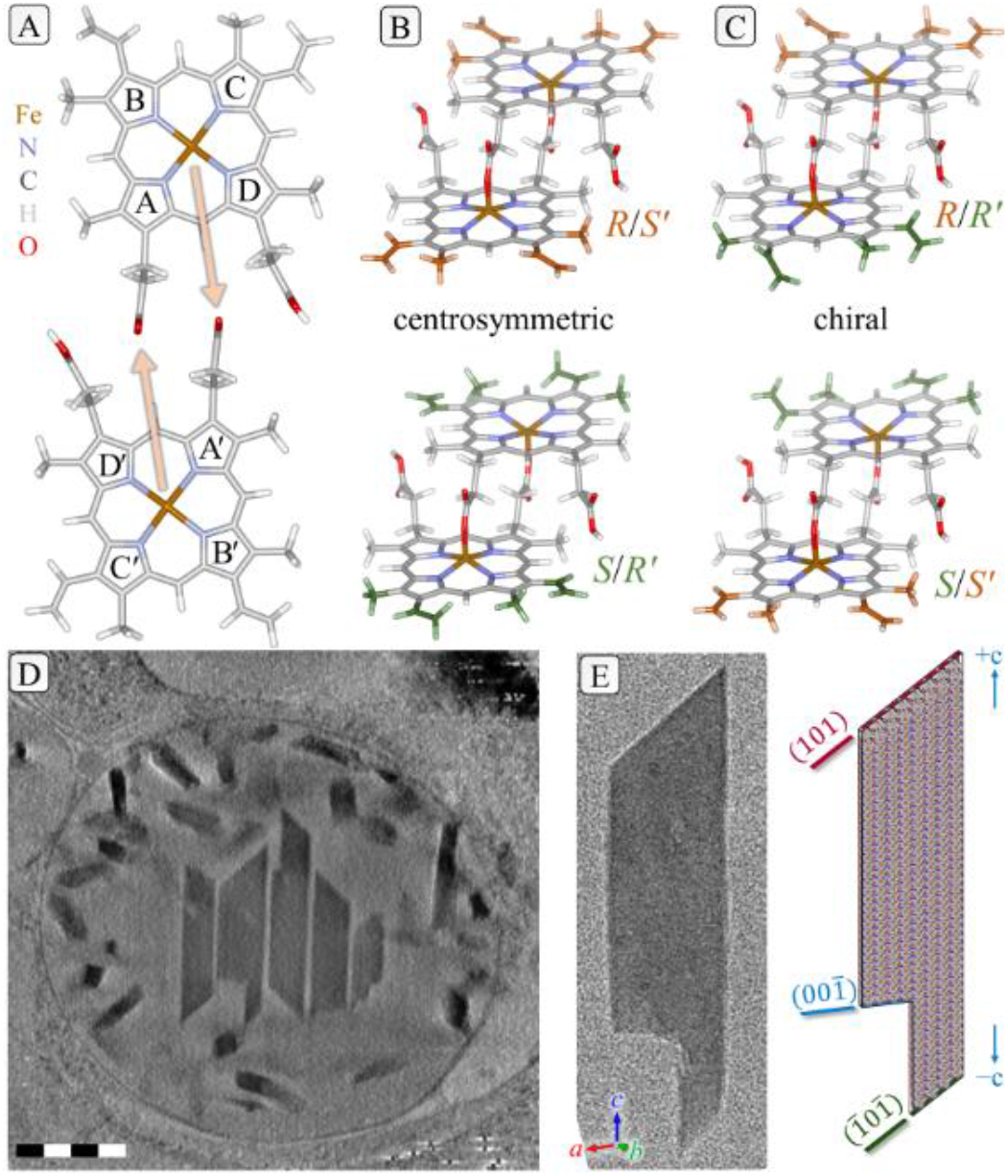
(A). Prior to crystallization, two heme monomers interlink to form cyclic hematin dimers via the binding of each central Fe ion to one of the two propionic acid tails (pyrrole ring labels included). The pro-chiral nature of the heme supports formation of four stereoisomers. (B). Centrosymmetric *R*/*S*′ and *S*/*R*′ dimers differ in binding of the Fe to the propionate moiety at the A or D pyrrole. (C). *R*/*R*′ and *S*/*S*′ are enantiomers. Colors of methyl and vinyl groups indicate the symmetrical relationship: both sides either orange or green for centrosymmetry and mixed for chiral dimers. See also Supporting Fig. S3. (D). Thick virtual section from a tomographic reconstruction of an intact digestive vacuole showing large crystals in the center and numerous smaller crystals decorating the periphery. Note the polar shapes of the crystals. The complete reconstruction appears in Supporting Movie S1. Scale bar 500 nm. (E). An exemplary crystal used in the diffraction study, with annotation of faces by *hkl* indices.

For the case of biogenic hemozoin, a chiral model of the hematin dimer was proposed by Straasø et al. on the assumption that the heme, upon release from oxygenated hemoglobin, retained the O_2_ molecule enantiotopically coordinated to the Fe(II) ion ^[11]^. An XRPD analysis of the crystal structure favored a 50:50 mixture of chiral (*S*/*S*′) and centrosymmetric (*R*/*S*′) dimers ^[11]^. However, the XRPD could not rule out an alternative interpretation.

Whole cell cryo-tomography using soft X-rays revealed biogenic hemozoin in the parasite digestive vacuole ^[18]^, and focused X-ray diffraction could suggest a preferential growth orientation ^[19]^. Recently developed cryogenic scanning transmission electron tomography (CSTET) provides a higher resolution view ^[20–22]^, while the development of 3D electron diffraction (3D ED) with narrow-field illumination makes it feasible to record diffraction patterns from sub-micron size specimens ^[23–26]^. In this work we enlist the new methods to address the crystal structure of biogenic hemozoin.

Two cell preparations were used in the present study. One was grown at 1% ambient O_2_ and the other at 5%, mimicking conditions for tissue and venous blood respectively. Infected red blood cells were deposited directly on electron microscope grids and then immediately vitrified. CSTET reconstructions of intact parasites revealed that opposite ends of hemozoin differ: one end shows a smooth chisel shape while the other is variable, typically squared off or ragged with an overhang (Fig. 1D). Similar shapes have been observed previously, either without comment ^[27,28]^ or with suggestion that the ragged end represents the site of growth ^[29]^. A purely centrosymmetric crystal structure should not produce such a polar morphology. Next, continuous-tilt diffraction data were recorded at −178 °C from isolated crystals that dispersed from ruptured cells. Indexing the crystal faces yielded the surprising result that the end cap orientations are not {011}, as seen in synthetic hemozoin, but rather (101) and 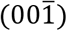, or (101) and a combination of 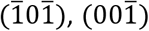, and nearby low index directions (Fig. 1E, Fig. S1). No significant differences were observed between the preparations under low and high oxygen atmospheres, and similar conclusions were reached from both. Details appear in Supporting Section 2.

A first complete set of 3992 Friedel-related *hkl* and 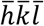reflections was obtained from 36 individual crystals grown at 1% O_2_. A second dataset from 64 crystals grown at 5% O_2_ yielded 8147 unique reflections using an improved detector system. In both cases the scaled intensities of corresponding reflections matched very well despite differences in crystal size, indicating the presence of only a single crystalline phase for biogenic hemozoin. Lattice parameters of biogenic hemozoin at low temperature were adopted from Straasø et al ^[11]^. Preliminary analysis and data reduction were performed using DIALS ^[30,31]^. Initial model structures of the three hematin dimers, *R*/*R*′, *R*/*S*′ and *S*/*R*′,were refined by least-squares electron structure factor analysis (Supporting Section 3) ^[32]^. The kinematic refinement cannot distinguish the chiral enantiomers, whereas the two centrosymmetric dimers differ in the coordination of the propionic acid on the *A* or *D* pyrrole to the neighboring Fe (Fig. 1). Given the polar crystal habits, these two dimers were not constrained as centrosymmetric entities. Overall *R* factor and atomic displacement parameters (ADPs) appear in Table S3. The fit of the *S*/*R*′ model was considerably worse than the others. The crystal structure embodying the chiral dimer *R*/*R*′ refined with fewest inconsistencies. All parameters refined well except the ADPs of terminal carbons in the vinyl substituents on the B′ and C′ pyrrole groups belonging specifically to the lower primed hematin ring. The related carbons on the upper monomer appeared sharp, however. A simple explanation is that the crystal structure contains a mixture of dimers, one chiral and one centrosymmetric (*R*/*S*′). This would account for the perfect atomic overlay on one side and the loss of atomic localization on the other for both the *R*/*R*′ and the *R*/*S*′ dimer. For roughly equal dimer occupancy, ADPs for the vinyl substituents were consistent with those of other constituent atoms. Additionally, all the vinyl groups assumed the more prevalent *cis* conformation, except for one that took a *trans* orientation. A further refinement based on dynamical diffraction analysis (Supporting Section 4) confirmed these conclusions as well as 1:1 occupancy of *R*/*S*′ and chiral dimers, as shown in Fig. 2 for the 5% O_2_ data and Fig. S5 for 1%.

**Figure 2.**
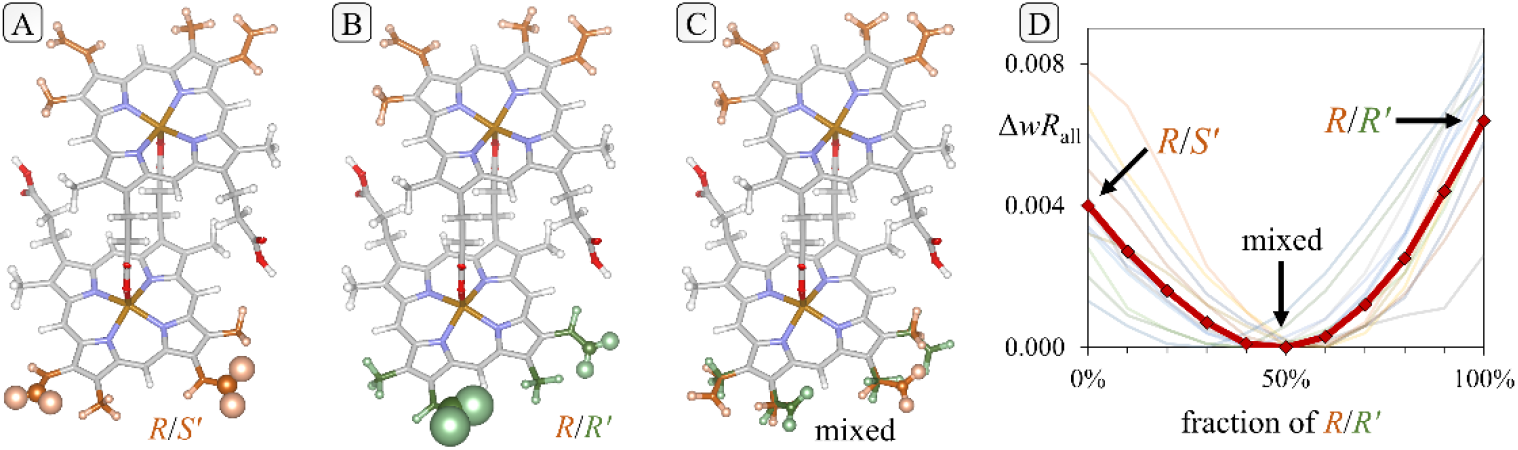
Refinement of the different models reveals a mix of chiral and centrosymmetric dimers. Atom spheres and ellipsoids represent displacement probability surfaces at the 10% level. (A). The centrosymmetric *R*/*S*′ model refines well except for the vinyl groups of a single monomer. (B). The chiral *R*/*R*′ model shows smaller ADPs and a single vinyl in the *trans* conformation. (C). All atoms refine well in a structure containing a mixture of *R*/*R*′ and *R*/*S*′ dimers. (D). R-factors of dynamical refinements with fixed composition indicate an equal concentration of *R*/*S’* and *R*/*R’* dimers in the structure. 13 faint lines represent the *R*-factor for a given composition based on the reflections from a single data set. The strong line is based on all reflections used in the refinement with a minimum close to 50% fraction of *R*/*R*′ dimers.

Density functional theory (DFT) calculations with the many-body dispersion (MBD) method ^[33]^ were performed to confirm: a) the plausibility of a crystal containing mixed isomers, and b) the *trans* orientation of one vinyl group belonging to a chiral dimer. Full unit cell relaxation was performed for crystals comprising pure and mixed dimers. Results are summarized in Fig. 3; details appear in Supporting Section 5. The outcome of DFT+MBD analysis, that the biogenic hemozoin crystal can easily comprise a mixture of *R*/*S*′ and one chiral dimer, supports the experimentally derived results. Other combinations have higher energy and are therefore less likely to occur. For the chiral dimer specifically, the vinyl on the C pyrrole is indeed in a *trans* conformation and the other three vinyl groups take a *cis* conformation. We attribute this to intermolecular forces in the crystal, which stabilize the otherwise less stable *trans* conformation.

**Figure 3.**
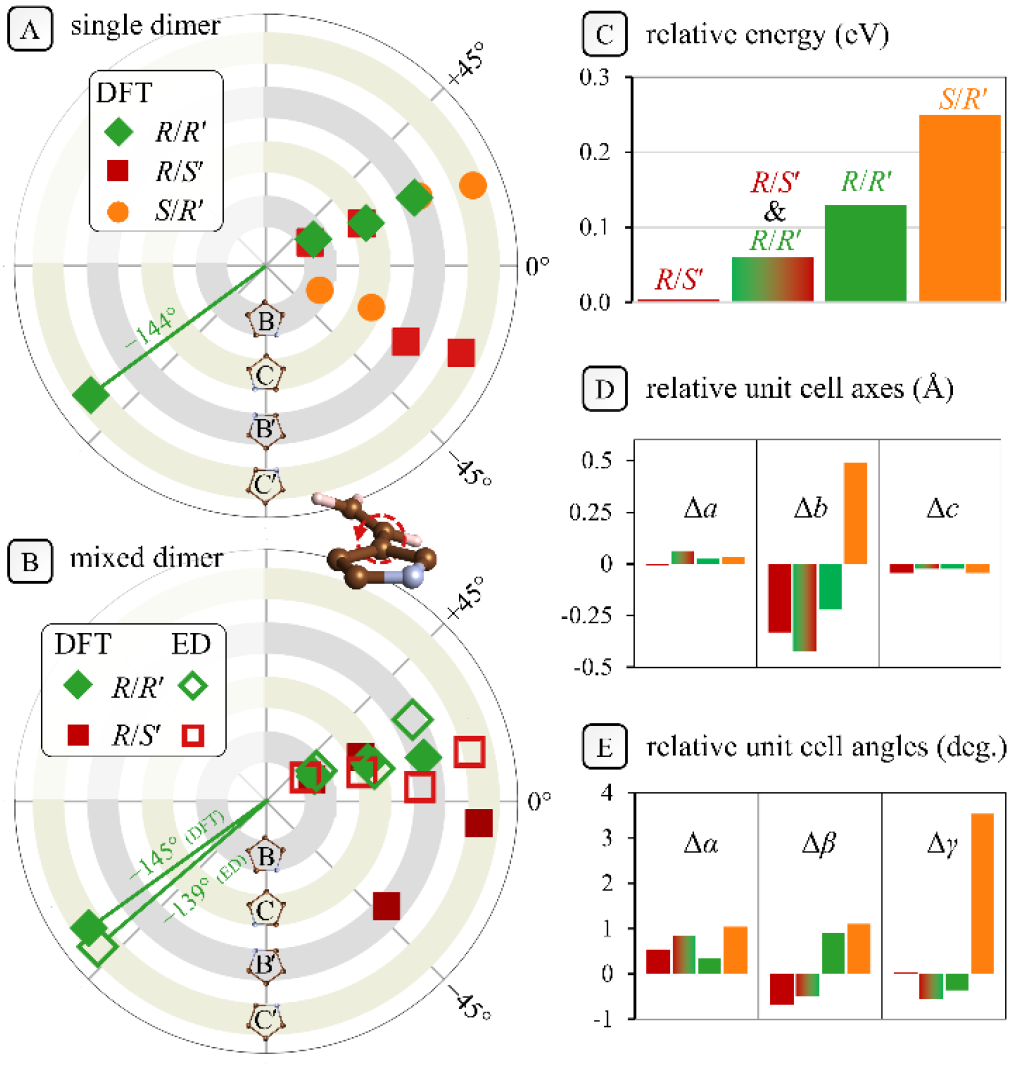
Results of DFT calculations. (A) All vinyl groups take the *cis* configuration, except for a single *trans* vinyl on the C′ pyrrole of the *R*/*R*′ dimer (see Fig. 1). (B) In the mixed *R*/*S*′+*R*/*R*′ crystal, again DFT predicts a single *trans* vinyl on the *R*/*R*′ dimer, in agreement with experiment (ED). (C) Energy rankings of the various models tested. (D,E) Unit cell parameters of the optimized structures in comparison with measurement ^[11]^. Color labels as in (C). Note the poor fit of the *S*/*R*′ dimer

Kinematic diffraction analysis cannot distinguish between the two chiral dimers. In principle, the interference of simultaneously excited beams within the crystal makes the diffracted intensities sensitive to the absolute structure ^[34]^. The effect is weak, however, given that mainly the terminal, half occupied vinyl CH_2_ groups contribute to the chirality, and we have been unable to discriminate the handedness with presently available methods (Tab. S5).

We may nonetheless deduce the absolute structure on the basis of morphology ^[35]^. A crystal composed purely of centrosymmetric *R*/*S*′ dimers should adopt a nonpolar habit, yet we observe a pronounced variability at one end. As seen in Fig. 1, the habit is polar, with a (101) face at the +*c* end and often a short 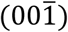face at the −*c* end; other crystals show a mixed shape at −*c* with a ragged 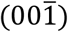 facet and a decidedly longer handle-shaped segment capped by a 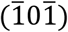facet (Figs. 1D and S1). Considering that the carboxyl pairs form H-bonded acid chains which run parallel to (101) and 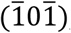, water in the aqueous medium of the digestive vacuole will bind poorly to either face. By comparison, the carboxyl and carboxylate groups at the 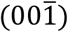 face are well exposed (Figs. 4 and S6-S8), so that growth of this face will be inhibited by competitive binding of water molecules. This explains why the habit of biogenic crystals differs from that of synthetic hemozoin grown in organic solvent (capped by {011} faces), and also why the 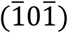 facet grows faster than 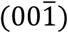 at the *–c* end. Considering the mixed crystal to be composed of *R*/*S*′ + *R*/*R*′ hematin dimers, atomic disorder will occur at the +b face and the –*c* end of the rod. Local inhomogeneities in composition could then explain the variable formation of the 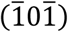 and 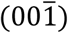 facets, reflecting centrosymmetric or polar structure (Fig. S1).

**Figure 4.**
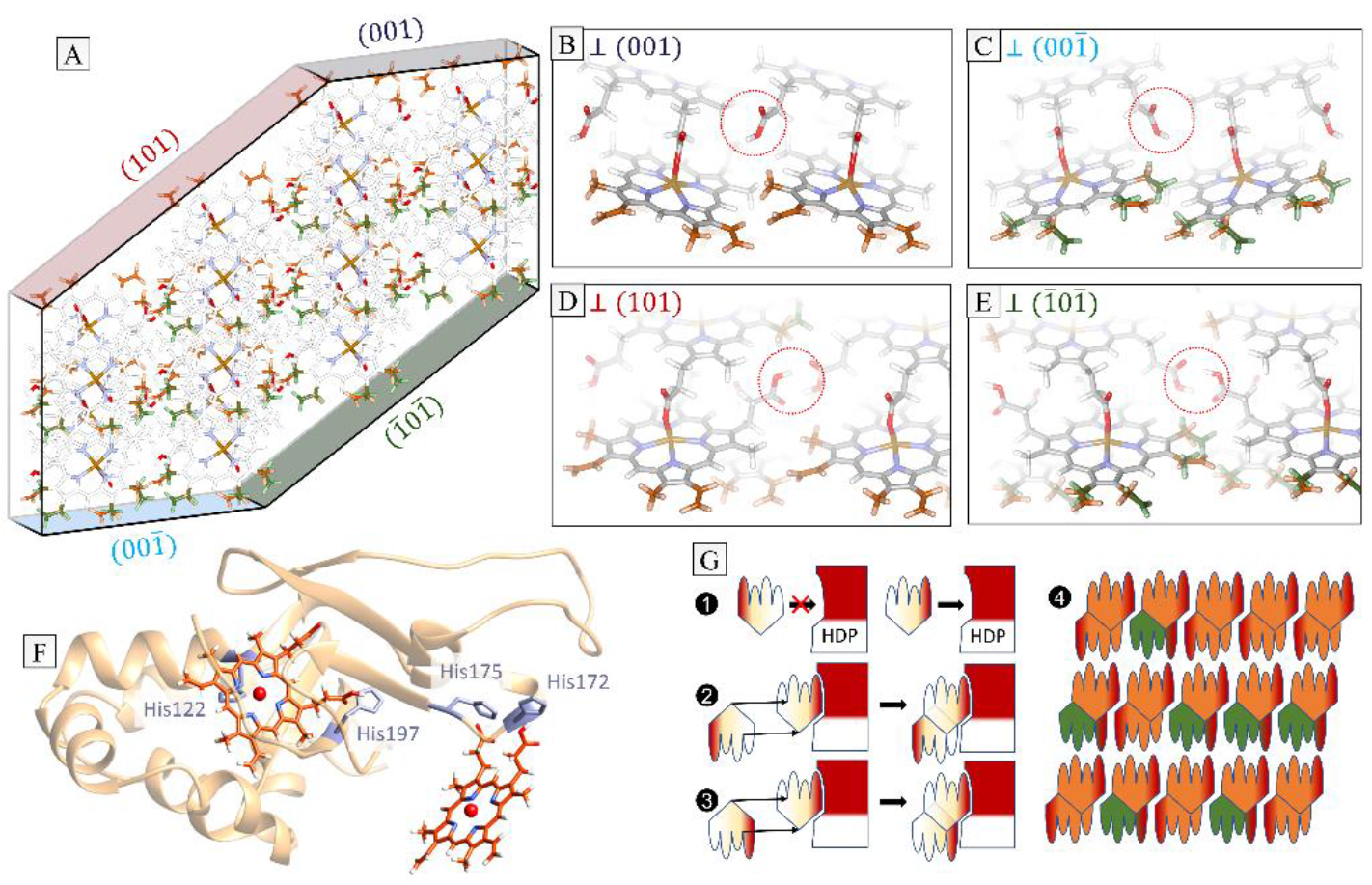
Groups exposed at the surfaces of hemozoin crystals composed of a mixture of *R*/*S*′ + *R*/*R*′ dimers. (A). Array of unit cells as visual guide. The mixture appears as disorder on the 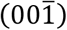 and 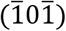 surfaces, but the (001) and (101) surfaces are completely ordered. (B-E). A detailed view of the methyl-vinyl pairs on the various exposed surfaces. Additionally, the carboxyl and carboxylate groups are exposed to the aqueous medium at the 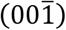 surface, rendering it more hydrophilic than the (101) and 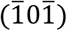 surfaces where the available bonds are saturated and buried (red circles, fading represents depth). Crystal growth of the relatively hydrophilic 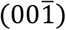 face will be inhibited by competitive binding of water. (F). A predicted structure for the heme detoxification protein (HDP) generated by AlphaFold ^[36]^ with heme groups docked manually to reflect histidine interactions ^[37]^. (G). A topological model for the function of HDP. A chiral agent that preferentially orients one monomer generates one chiral and one centrosymmetric dimer, resulting in a polar crystal where disorder appears on only one side.

Morphological analysis of biogenic hemozoin recalls the assignment of molecular chirality in centrosymmetric crystals composed of a mixture of chiral *R* and *S* molecules grown in the presence of a tailor-made chiral inhibitor (*R*′ or *S*′) ^[35]^. In this regard, we present here the first reported case of such assignment of handedness for a molecule of unknown chirality.

In another approach, we performed through-focus imaging and phase retrieval ^[38]^ on a crystal in a favorable orientation along the −***a*** direction, where the vinyl-methyl groups are visible in otherwise empty channels. Multi-slice simulations using the refined models generated in the dynamical diffraction analysis yielded a superior fit to the data for the mixture containing the *R*/*R*′ rather than the *S*/*S*′ chiral dimer (Supporting Section 6).

The selection of one chiral and one centrosymmetric dimer implies the presence of a chiral agent in their formation. The heme detoxification protein (HDP) is associated with falcipain 2 and other proteases, and has been implicated as a catalyst for dimerization between steps of hemoglobin digestion and hemozoin crystallization ^[39]^. Biochemical studies identified four key histidine residues: His122 and His197 form one heme binding site, with His122 as the ligand to iron, while His172 and His175 form another ^[37,40]^. To date there is no high-resolution molecular structure available for HDP, although it is recognized to have a fascilin-like fold. Therefore, we turned to structural modeling by AlphaFold^[36]^ and other servers (Fig. 4F&G and Supporting Section 7). There was a broad consensus at high confidence in the architectural features, with a compact α-helical domain at the N terminus and a largely unfolded cap at the C terminus. One heme binding site containing the Fe-binding histidine appears in a pocket between the compact core of the protein and the cap, while the other appears at the tip of a long unfolded loop. The dimerization reaction will occur between these two His-bound hemes. If chiral discrimination at the protein core fixes the orientation of one heme, then topologically, only one chiral dimer may form (Fig. 3G, Fig. S12).

Heme is oxygenated on the *R* face in hemoglobin. If heme remains oxygenated after degradation of hemoglobin, a chiral S/S’ dimer might form spontaneously, as proposed in an earlier model ^[11]^. Intermediate binding of oxygenated heme to HDP via His122 would reverse this orientation, resulting in the *R*/*R*′ dimer (See Fig. S9 in ^[17]^). According to the oxygen-binding properties of hemoglobin, 1% ambient O_2_ corresponds to heme oxygenation below 10%, whereas 5% O_2_ provides a saturation of 75%. In both cases the diffraction data indicated a similar concentration of about 50% chiral dimer, suggesting that the dimer stoichiometry is under biological rather than chemical control. Such regulation would not be possible for synthetic crystals.

In summary, tomography, diffraction, density functional theory, morphological analysis and through-focus imaging combine to yield a refined structure of native hemozoin from Plasmodium falciparum. The crystal structure contains a 1:1 mixture of hematin dimers, one centrosymmetric and one chiral, and previously unrecognized, deeply corrugated (101) and 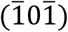 surfaces. This definitive structure should serve as a template for development of treatments for malaria where hemozoin is a drug target.

## Acknowledgements

We thank Jens Als Nielsen for a critical reading of the manuscript and Isabella Weissbuch for inspiring discussions. This work was funded in part by grants to ME from the Minerva Foundation, the Israel Science Foundation (project 1696/18), and the Estate of David Levinson, to LP from the Czech Science Foundation (project 21-05926X) and the Czech Ministry of Education, Youth and Sports, LM2018110 (CzechNanoLab Research Infrastructure). ME is Head of the Irving and Cherna Moskowitz Center for Nano and Bionano Imaging and incumbent of the Sam and Ayala Zacks Professorial Chair in Chemistry. We acknowledge Diamond for access and support of the cryoEM facilities at the U.K. national electron Bio-Imaging Centre (eBIC) (proposals NT21004 and NT29812), funded by the Wellcome Trust, MRC, and BBSRC. Work at CMU was supported by the National Science Foundation (NSF) Division of Materials Research (DMR) through grant DMR-2131944. This research used computational resources of the National Energy Research Scientific Computing Center (NERSC), a DOE Office of Science User Facility supported by the Office of Science of the U.S. Department of Energy, under Contract DE-AC02-05CH11231; the Argonne Leadership Computing Facility (ALCF), which is a DOE Office of Science User Facility supported under Contract DE-AC02-06CH11357; and the Extreme Science and Engineering Discovery Environment (XSEDE), which is supported by National Science Foundation grant number ACI-1548562. This work was supported in part by the Wellcome Trust Investigator Award (206422/Z/17/Z) and the ERC AdG grant (101021133).

For the purpose of open access, the authors have applied a Creative Commons Attribution (CC BY) licence to any Author Accepted Manuscript version arising.

All data needed to evaluate the conclusions in the paper are present in the paper and/or the Supporting Information. Diffraction datasets are published at https://doi.org/10.5281/zenodo.5039355 and https://doi.org/10.5281/zenodo.7462145.

## Notes

https://doi.org/10.5281/zenodo.5039355

https://doi.org/10.5281/zenodo.7462145

### Competing Interest Statement

The authors have declared no competing interest.

### Summary of Updates

addition of hemozoin diffraction data from cells grown under 5% oxygen.

