## Supplementary Information for "Polar Crystal Habit and 3D Electron Diffraction Reveal the Malaria Pigment Hemozoin as a Selective Mixture of Centrosymmetric and Chiral Stereoisomers"

#### 1. Experimental Methods

##### Specimen Preparation

Two cultures of red blood cells (RBC) infected by Plasmodium falciparum (NF54) were prepared independently under 1% (batch 1) and 5% (batch 2) oxygen partial pressure, and separated magnetically. Cells were fixed with 0.5% glutaraldehyde in PBS for 5 min and washed three times with serum-free medium. Cells were then applied to standard Quantifoil EM grids having round holes of 3.5 µm diameter, blotted with filter paper from behind in a Leica EM-GP automated instrument, and plunged into liquid ethane for vitrification. As the hole size is marginally smaller than the RBC, the cells tend to lodge and burst, exposing the parasites. Often the parasites also burst, spreading the contents, including hemozoin, onto the surrounding area. The time interval from blotting to plunging is at most a few seconds.

##### Cryo-Scanning Transmission Electron Tomography (CSTET)

STEM tomography data of samples from batch 1 were acquired on a Talos Arctica or Titan Krios microscope (Thermo Fisher Scientific) operating at 200 or 300 kV acceleration using the manufacturer's software. Typical STEM parameters were: semi-convergence angle 1.2 mrad, probe current 20 pA, pixel sampling 2 nm/pixel, and scan frame time 50 sec. Tilt angles spanned −60 to 60 deg in two sweeps. On the Talos, the bright field detector was used with a camera length of 330 mm, equivalent to a collection semi-angle of 3 mrad. On the Krios, an incoherent brightfield signal was collected using the Fischione HAADF detector in combination with a 20 µm objective aperture, yielding a collection semi-angle of approximately 3 mrad. Reconstructions were made using IMOD [1] by back-projection with the SIRT-like filter (simultaneous iterative reconstruction technique).

##### Electron diffraction data collection

TEM diffraction data of batch 1 were collected using a Talos Arctica (Thermo Fisher Scientific) operated at 200 kV with a stage temperature of ~80 K. The optical setup aimed to minimise electron dose whilst also reducing exposure to crystals in the immediate area. To achieve a minimal current, the largest gun lens setting (8) and the largest spot size (11) were used with a C2 aperture of 50 µm. The beam was condensed to 1.78 µm diameter. The diffraction patterns were collected at a nominal camera length of 1.05 m. The dose rate under these conditions was 0.04 e $Å^{-2}\,s^{-1}$. A Ceta-D detector (Thermo Fisher Scientific) was used for diffraction data collection. The goniometer stage and detector were controlled by EPU-D software (Thermo Fisher Scientific). Crystals were initially tilted from +48° to +46° to correct for backlash followed by 1° of continuous rotation prior to collection starting at +45°. All data sets covered 90° of goniometer rotation with a collection speed of 1° or 0.5° per second. The exposure time for each frame was 1 second. Each frame was written as an mrc file and stacked using IMOD [1]. The diffraction data sets of batch 1 are published at https://doi.org/10.5281/zenodo.5039355.

TEM diffraction data of batch 2 were collected using a Glacios microscope (Thermo Fisher Scientific) operated at 200 kV with a stage temperature of ~80 K and similar beam settings to those described above for the Arctica. The diffraction images were recorded by a DECTRIS SINGLA detector at a nominal camera length of 677 mm according to the microscope control software (estimated to be to 809 mm at the detector). The sample was rotated through 90° over a 18 seconds with the camera collecting data at 50 Hz, producing images representing 0.1°. Subsequently, the goniometer rotation angle was refined to 0.11° per frame. The dose rate under these conditions was 0.01 $e^{-}$ $Å^{-2}$ $s^{-1}$. These diffraction data sets of batch 2 are published at https://doi.org/10.5281/zenodo.7462145.

In order to examine the relation between the polar shape and the axis orientations, images of a collection of hemozoin crystals from batch 1 were collected on the Talos Arctica microscope with Falcon III detector in linear mode at 36000x magnification (calibrated pixel size 4.1 Å/px). Diffraction data were also collected from these crystals to determine the orientation of the unit cell and thus index the crystal faces.

#### 2. Crystal Morphology and Indexing

Morphological analysis consisted of measurements of two lengths along either side of these crystals using ImageJ [2]. A transverse width was calculated between these two lines, and assigned to be parallel to either the *a** or *b** direction according to the determination of the orientation matrix. The results are given in Table S1 and an example shown graphically in Figure S1.

From these results we observe that the "chisel" end lies in the +c orientation $(001)$ while the "ragged" end lies in the –c orientation $(00\bar{1})$. The "chisel" face is in the $(101)$ orientation rather than $(011)$ as reported for synthetic hemozoin, while the "ragged" end exposes variable lengths of low-index faces including $(100)$, $(\bar{1}00)$, $(010)$, $(0\bar{1}0)$, and notably $(00\bar{1})$, $(\bar{1}0\bar{1})$.

| Data set | Crystal | L1 (nm) | L2 (nm) | W (nm) | Aspect ratio max(L1,L2)/W | Beam direction |
|---|---|---|---|---|---|---|
| 2 | 1 | 498.3 | 525.0 | 79.3 | 6.6 | +a |
| | 2 | 453.5 | 446.7 | 80.6 | 5.6 | +b |
| | 3 | 442.4 | 533.5 | 88.2 | 6.1 | +b |
| 3 | 1 | 463.7 | 522.4 | 108.9 | 4.8 | +b |
| 4 | 1 | 543.8 | 516.5 | 84.9 | 6.4 | −a |
| 5 | 1 | 251.8 | 286.7 | 94.4 | 3.1 | +b |
| | 2 | 534.1 | 412.1 | 118.9 | 4.5 | −b |
| | 3 | 559.7 | 458.4 | 114.2 | 4.9 | +b |
| 7 | 1 | 460.8 | 393.3 | 113.0 | 4.1 | −a |
| 8 | 1 | 421.3 | 257.5 | 146.3 | 2.9 | −b |
| 10 | 1 | 417.0 | 436.7 | 166.0 | 2.6 | +b |
| 11 | 1 | 617.5 | 686.4 | 210.6 | 3.3 | +a |
| | 2 | 757.7 | 513.2 | 193.1 | 3.9 | +b |
| 12 | 1 | 618.7 | 934.4 | 250.7 | 3.7 | −b |
| 13 | 1 | 499.6 | 791.0 | 212.8 | 3.7 | −b |
| | 2 | 773.0 | 562.1 | 239.8 | 3.2 | −a |
| 14 | 1 | 461.1 | 425.9 | 165.0 | 2.8 | −b |
| 15 | 1 | 303.6 | 295.6 | 78.0 | 3.9 | +b |
| 16 | 1 | 656.6 | 579.1 | 109.9 | 6.0 | +b |

**Table S1.** Crystal size measurements performed on real space images, including the assignment of the transverse axis by comparison with diffraction data indexing. Beam direction indicates which unit cell axis is approximately parallel (minus sign means antiparallel) to the primary electron beam at a goniometer angle of 0°. The data sets are not sequentially numbered because 1, 6, and 9 were discarded during analysis when individual crystals could not be indexed and discerned in shape as well.

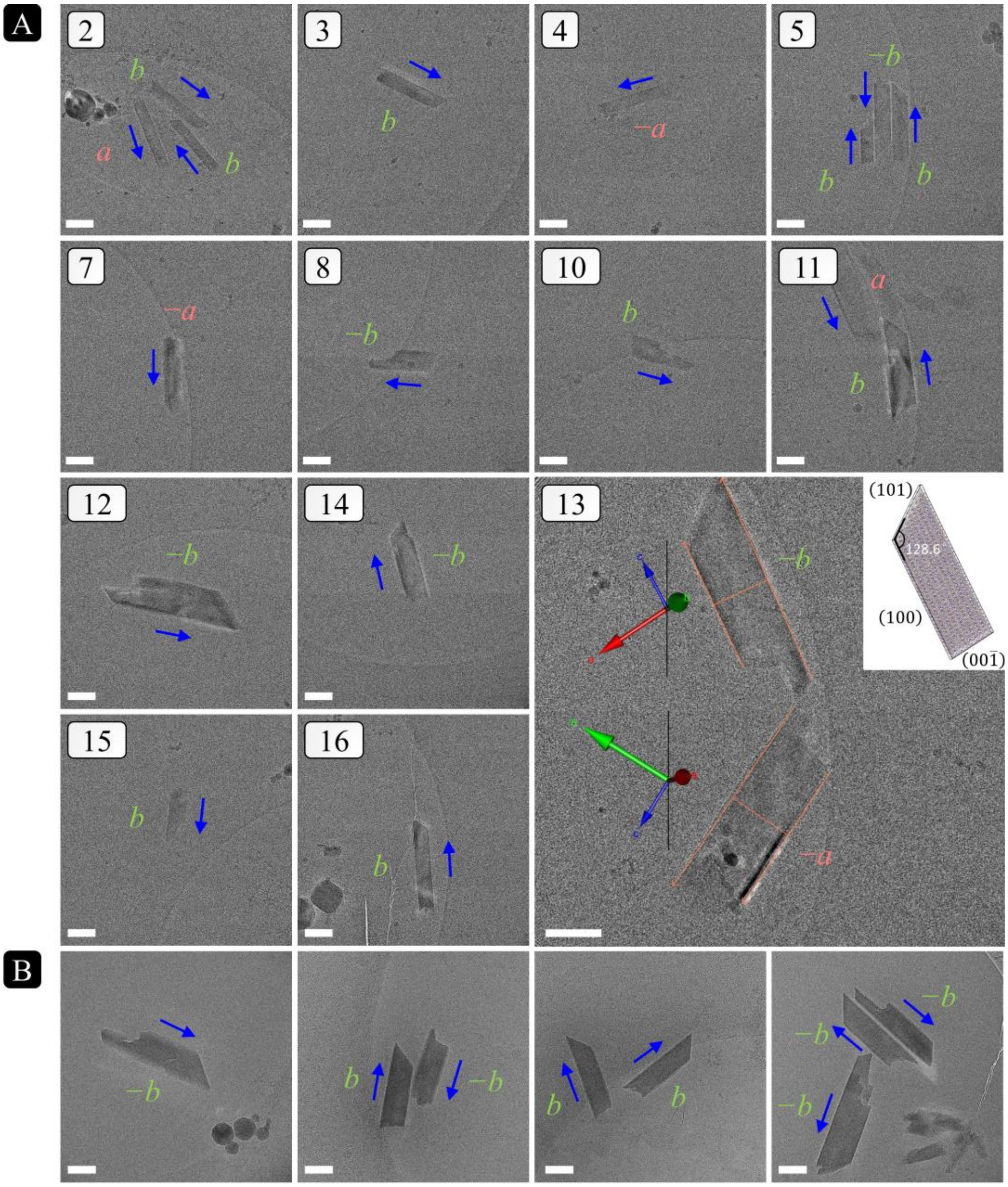

**Figure S1.** Real-space images of hemozoin crystals. The white scale bar corresponds to a length of 200 nm. Blue arrows indicate the direction of the c-axis. Assignment of the axes was performed by comparison with the indexed lattice from diffraction. The letter next to each crystal is the unit cell axis closest to the viewing direction (parallel to the electron beam). (A) Micrographs corresponding to data sets of batch 1 of Table S1, using the same numbering. Image of data set 13 shows the two crystals with orange lines overlaid used to measure their lengths (along the *c* axis) and transverse widths (along either *a** or *b**). The inset shows a model crystal with the orientation of the upper crystal and indices of selected faces. (B) Images from four representative images from batch 2 used for the structural analysis.

#### 3. Structure analysis and kinematical refinement

##### Data processing

Electron diffraction data of batch 1 were recorded in MRC format, either as individual images or image stacks, using the Ceta-D camera and EPU-D software. These were read into DIALS 3.5.0, which can interpret the MRC format directly without conversion. The known tilt range per image, effective detector distance and beam centre were provided as additional metadata during import. As seen previously, a negative bias was evident in the background intensities of images from the Ceta-D detector [3]. We investigated a range of additive correction factors to account for the bias, but an attempt to find the optimum level was inconclusive. The final structures were comparable, for the purpose of the conclusions drawn here, across the processing runs with different pedestal settings, and model quality was not significantly better than adding no pedestal to correct for the bias. For that reason, we omitted the bias correction for all data sets of batch 1 here.

In some cases, a few images at the end of a tilt series were excluded from processing, where these showed poor diffraction quality. A backstop mask was provided to exclude the shadowed regions from spot-finding and integration. Multiple diffracting crystals were evident in 10 out of 19 tilt series ultimately used for structure determination. These tilt series were recorded from 17 independent positions, with repeat tilt series recorded at two positions. A total of 38 lattices were successfully indexed from these 19 tilt series from batch 1.

Diffraction data from crystals of batch 2 were recorded in HDF5 format using the SINGLA detector (DECTRIS). These were read by DIALS 3.12.0, with missing metadata supplied during import. The SINGLA is an electron-counting detector, and we assumed a count multiplicity or detector gain of exactly 1. No backstop was used during data collection. Spurious reflections were observed at the edges and corners of the detector, presumably caused by scattering at the mounting flange. We provided a mask to exclude regions of the detector that would capture reflections beyond 0.7 Å resolution.

We included data collected from 32 tilt series. Multiple lattices were evident from at least 18 tilt series. In total, we included data from 64 separately indexed lattices.

For each tilt series of both batches, geometry refinement proceeded in stages for stability. The effective detector distance was fixed during initial refinement after indexing. A subsequent round of geometry refinement was then performed, allowing the detector parameters to vary, but applying a restraint to the unit cell $a$ = 12.086 Å, $b$ = 14.6216 Å, $c$ = 7.9942 Å, $\alpha$ = 90.758°, $\beta$ = 97.093°, $\gamma$ = 97.060° previously determined by X-ray powder diffraction at 80 K [4]. During the final rounds of refinement, the effective detector distance was again fixed, while the overall cell parameters were allowed to refine freely and a smoothly-varying model for the crystal orientation was determined [5]. The model for each experiment thus determined was used to integrate reflections from 38 (batch 1) and 64 (batch 2) indidivual crystals.

For each batch the integrated data sets were combined and scaled by dials.scale, making use of automated image group exclusion by the $\Delta CC_{1/2}$ method [6]. Data processing statistics for the combined data sets are shown in Table S2. Despite the *P*1 symmetry and limited tilt range for each individual data set, the combined reflections are 100% complete to a resolution of 0.90 Å (batch 1) and 0.71 Å (batch 2). For batch 1, the scaled reflections were written to unmerged MTZ format before conversion to hkl format using mtz2hkl. For batch 2, the hkl file was written out directly by DIALS 3.12.0. In both cases, the unit cell was set to the above referenced parameters from powder diffraction.

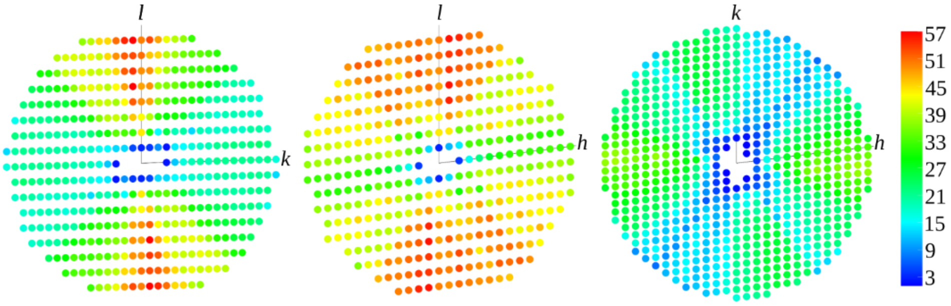

**Figure S2.** Multiplicity of reflections in slices (0*kl*), (*h*0*l*) and (*hk*0) adapted from the output of cctbx.multiplicity_viewer for the combined data set from batch 1.

| | Batch 1 | | | Batch 2 | | |
|---|---|---|---|---|---|---|
| | Overall | Low-resolution shell | High-resolution shell | Overall | Low-resolution shell | High-resolution shell |
| High-resolution limit (Å) | 0.90 | 2.43 | 0.90 | 0.71 | 1.93 | 0.71 |
| Low-resolution limit (Å) | 9.82 | 9.82 | 0.92 | 14.51 | 14.51 | 0.72 |
| Completeness (%) | 100.0 | 99.5 | 100.0 | 100.0 | 100.0 | 100.00 |
| Multiplicity | 31.7 | 26.1 | 28.9 | 57.5 | 50.5 | 61.0 |
| $I/\sigma$ | 8.2 | 31.2 | 1.3 | 6.7 | 33.2 | 0.8 |
| $R_{meas}(I)$ | 0.344 | 0.126 | 4.146 | 0.592 | 0.194 | 30.976 |
| $R_{pim}(I)$ | 0.063 | 0.026 | 0.820 | 0.078 | 0.028 | 3.946 |
| $CC_{1/2}$ | 0.999 | 0.999 | 0.642 | 0.997 | 0.998 | 0.529 |
| $R_{Friedel}(F)$ [7] | 0.084 | | | 0.063 | | |
| Total observations | 126299 | 5148 | 5799 | 468746 | 20644 | 26727 |
| Total unique | 3990 | 197 | 201 | 8147 | 409 | 438 |

**Table S2.** Data processing statistics. $R_{Friedel}(F)$ calculated as described in [7]

#### Model refinement

The following steps are valid for both batches as during structure solution and refinement no significant differences were observed between the analysis of the two scaled data sets. A SHELX INS file was generated by edtools.make_shelx [8], which includes electron scattering factors. Initial structural models in space groups $P\bar{1}$ and $P1$ were determined by SHELXT [9]. The $P\bar{1}$ solution was taken as the start point for the construction of additional models, which were refined with SHELXL version 2019/2 [10]. Electron scattering factors tabulated in Table 4.3.1.1 of [11] were fitted against the Cromer-Mann-parametrisation with GNUPLOT [12]. Starting values for the fitting were taken from the parametrisation in [13].

##### *Reference coordinate system*

When referring to our models, we place the origin of the unit cell at the top left corner. The *a*-axis runs horizontally to the right, the *b*-axis runs vertically down (Fig. S3). In this orientation, the *c*-axis runs horizontally backwards away from the viewer. This completes a right-handed coordinate system. In this orientation, the hemozoin dimer is composed of an upper and a lower hematin unit. The *y*-coordinate of the central Fe-ion of the upper hematin is less than the *y*-coordinate of the central Fe-ion of the lower hematin.

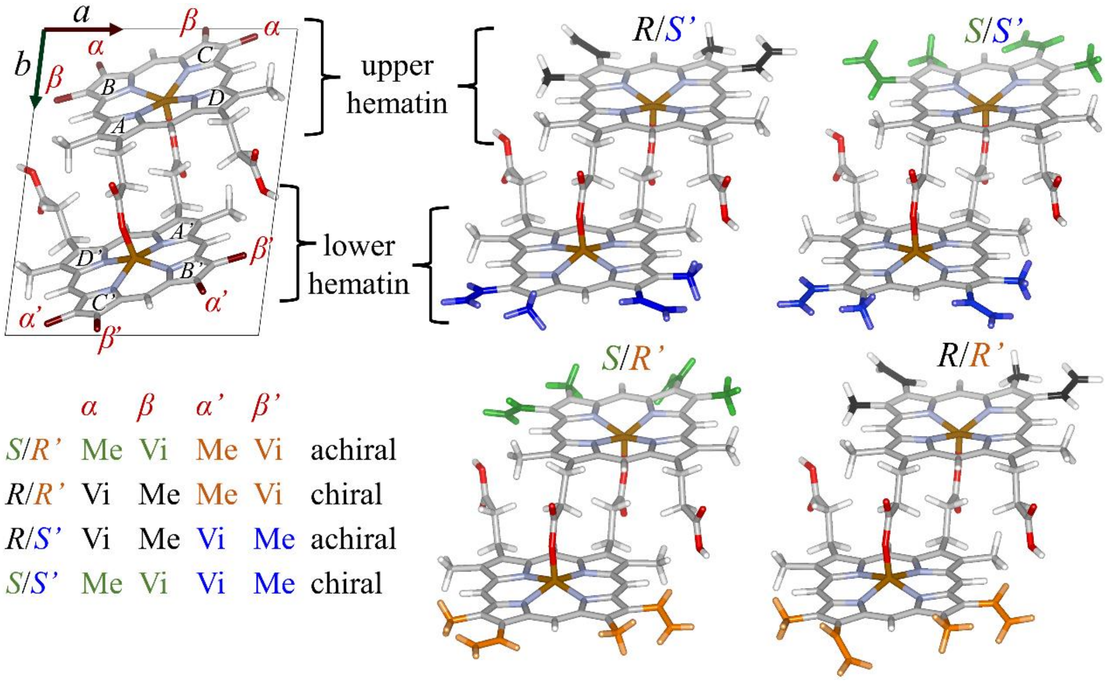

| | α | β | α' | β' | |
|---|---|---|---|---|---|
| *S*/*R'* | Me | Vi | Me | Vi | achiral |
| *R*/*R'* | Vi | Me | Me | Vi | chiral |
| *R*/*S'* | Vi | Me | Vi | Me | achiral |
| *S*/*S'* | Me | Vi | Vi | Me | chiral |

**Figure S3.** Orientation and position of hematin anhydride dimer in the unit cell. Labels of pyrrole rings (A to D, A′ to D′) and relevant sites for methyl (Me) and vinyl (Vi) groups (α, β, α′, β′) included. Figure inspired by Straasø et al. (2014). Here color indicates the stereochemical label of the upper (*R* = black, *S* = green) and lower (*R*′ = orange, *S*′ = blue) hematin monomer. This complements the color scheme of Figure 1 in the main text where color denotes the symmetrical relationship.

*Models*

We refer to *R*-hematin and *S*-hematin as described in [4] and used in the main text. To distinguish both moieties, we prime the lower hematin (Fig. S3).

The hematin anhydride dimers *R*/*S*′ and *S*/*R*′ are achiral. Respective models were refined in the centrosymmetric space group $P\bar{1}$ as well as in the space group $P1$. In $P\bar{1}$ the coordinates and atomic displacement parameters (APDs) of the upper and lower hematin are constrained by symmetry to be pairwise equal. Refinements of *R*/*S*′ and *S*/*R*′ in $P1$ start from the same configuration, but during refinement, the upper and lower hematin are not constrained to one another. Note that we did apply geometric restraints generated with the GRADE server (http://grade.globalphasing.org/cgi-bin/grade/server.cgi) [14]. For disordered models with two partially occupied dimers in the unit cell, only the enantiotopic methyl and vinyl groups at the indicated sites (Table S3) were modelled as superposed disorder, which is described in more detail below. *R*/*R*′ and *S*/*S*′ are enantiomers, which cannot be distinguished within the kinematical approximation (see Section 4 of this document). Due to reasons given in the main text, we here exclude models involving *S*/*S*′ hematin anhydride dimers.

Finally, we also carried out a rigid body refinement of the superposition of *R*/*S*′ and *R*/*R*′, i.e. the model of the *R*/*S*′ dimer was allowed to translate and rotate as one entity, and the model of the *R*/*R*′ dimer was allowed separately to translate and rotate.

| Modelled dimers | Disorder sites | Space group | occ. #1 | No. of params | $R_{obs}$ | $R_{all}$ | $wR2_{all}$ | ⌀ $U_{iso}$ (Å²) | max. $U_{iso}$ (Å²) |
|---|---|---|---|---|---|---|---|---|---|
| **Batch 1** | | | | | | | | | |
| **Ordered single-dimer refinements** | | | | | | | | | |
| *S*/*R*′ | | $P\bar{1}$ | 1.0 | 177 | 0.217 | 0.239 | 0.525 | 0.075 | 0.59(7) |
| *R*/*S*′ | | $P\bar{1}$ | 1.0 | 177 | 0.204 | 0.227 | 0.493 | 0.061 | 0.221(13) |
| *S*/*R*′ | | *P*1 | 1.0 | 350 | 0.187 | 0.208 | 0.459 | 0.098 | 2.0(9) |
| *R*/*S*′ | | *P*1 | 1.0 | 353 | 0.174 | 0.195 | 0.429 | 0.081 | 1.2(3) |
| *R*/*R*′ | | *P*1 | 1.0 | 353 | 0.170 | 0.191 | 0.426 | 0.066 | 0.59(10) |
| **Refinements with disordered methyl and vinyl groups** | | | | | | | | | |
| *S*/*R*′ & *R*/*R*′ | α, β | *P*1 | 0.231 | 842 | 0.142 | 0.163 | 0.365 | 0.056 | 0.129(8) |
| *R*/*S*′ & *R*/*R*′ | α′, β′ | *P*1 | 0.50 | 840 | 0.137 | 0.158 | 0.344 | 0.058 | 0.109(7) |
| *R*/*S*′ & *S*/*R*′ | α, β, α′, β′ | *P*1 | 0.62 | 899 | 0.135 | 0.157 | 0.342 | 0.058 | 0.109(7) |
| **Batch 2** | | | | | | | | | |
| **Ordered single-dimer refinements** | | | | | | | | | |
| *S*/*R*′ | | $P\bar{1}$ | 1.0 | 392 | 0.175 | 0.228 | 0.461 | 0.049 | 0.165(6) |
| *R*/*S*′ | | $P\bar{1}$ | 1.0 | 392 | 0.157 | 0.212 | 0.421 | 0.049 | 0.131(4) |
| *S*/*R*′ | | *P*1 | 1.0 | 350 | 0.177 | 0.242 | 0.443 | 0.050 | 0.239(18) |
| *R*/*S*′ | | *P*1 | 1.0 | 353 | 0.163 | 0.231 | 0.418 | 0.046 | 0.231(17) |
| *R*/*R*′ | | *P*1 | 1.0 | 353 | 0.160 | 0.228 | 0.414 | 0.044 | 0.183(13) |
| **Refinements with disordered methyl and vinyl groups** | | | | | | | | | |
| *S*/*R*′ & *R*/*R*′ | α, β | *P*1 | 0.753 | 382 | 0.160 | 0.228 | 0.411 | 0.045 | 0.197(14) |
| *R*/*S*′ & *R*/*R*′ | α′, β′ | *P*1 | 0.543 | 380 | 0.160 | 0.228 | 0.404 | 0.043 | 0.147(11) |

**Table S3.** Ordered (one dimer) and disordered (two dimers) models of hemozoin refined applying the kinematical theory of diffraction. Occ. #1 is the refined constrained occupancy of the first dimer. ⌀ $U_{iso}$ and max. $U_{iso}$ are the average and maximum (equivalent) isotropic displacement parameters of non-hydrogen atoms, respectively.

##### *Restraints used during refinement*

Hydrogen atoms were refined in riding positions with internuclear bond distances.

The initial solution was converted to a PDB file with the command 'WPDB -1' in SHELXL. The PDB file was reduced to a single moiety and converted to mol2 format with openBabel [15]. Geometric restraints were calculated with the GRADE server [14] from this mol2-file.

ADPs were sometimes refined isotropically and restrained with the SHELXL command "SIMU_* 0.01 0.02". We say 'sometimes', because for the interpretation of the model as isomeric disorder, restraints of the ADPs were turned off. This resulted in unusually high ADP values only for one of the two hematin monomers. In the *R*/*R*′ model, for example, both terminal C-atoms of the two vinyl groups at the β′ sites assumed very large ADP values, while the ADPs of all carbon atoms of the upper hematin (α and β site) assumed reasonable values. We interpreted this as an indication of isomeric disorder of one hematin.

In the refinements of non-disordered single-dimer models, BUMP restraints were used and had a strong impact on the dihedral angles of the methy-vinyl groups. In the presence of BUMP, several dihedral angles were squashed to values near 0°. This results in a van-der-Waals clash with the nearest methyl carbon. Presumably, the anti-bumping from the packing had a stronger impact than this van-der-Waals clash. This might indicate incorrect unit cell parameters. However, an optimization of the unit cell parameters against geometric restraints with  the tool CELLOPT [16] did not resolve the van-der-Waals clash in the refined single-dimer models. This is another indication for the presence of isomeric disorder.

##### *Scan of Dihedral Angles*

The following procedure was followed in order to avoid local minima of the dihedral angles of each methyl-vinyl group. For every model the Cartesian coordinates of the atoms C14, C15, C19, C20 were copied from the PDB file. The starting configuration had an angle of 6°. The following OCTAVE script was used to rotate the C20-atom about the C15-C19 bond in steps of 45°:

```
#################### start script ###########################################
# ortho coords WPDB 1
C20 = [ 9.343 ; -0.623 ;  9.766 ]
C19 = [ 9.000 ; 0.249 ; 8.820 ]
C15 = [ 7.650 ; 0.766 ; 8.614 ]
C14 = [ 6.539 ; 0.630 ; 9.384 ]

axis = C19 - C15

uaxis = axis/norm(axis)

x = uaxis(1)
y = uaxis(2)
z = uaxis(3)

angle = 45

phi = pi/180.*angle

M = [ x*x*(1-cos(phi)) + cos(phi), x*y*(1-cos(phi)) - z*sin(phi), x*z*(1-
cos(phi)) + y*sin(phi);
      x*y*(1-cos(phi)) + z*sin(phi), y*y*(1-cos(phi)) +   cos(phi),  y*z*(1-
cos(phi)) - x*sin(phi);
      x*z*(1-cos(phi)) - y*sin(phi), y*z*(1-cos(phi)) + x*sin(phi), z*z*(1-
cos(phi)) +   cos(phi) ]

printf ("angle = %4.1f: %8.3f%8.3f%8.3f\n", 0, C20(1), C20(2), C20(3))
for i = 1:8
C20 = M*(C20 - C19) + C19;
printf ("angle = %4.1f: %8.3f%8.3f%8.3f\n", i*angle, C20(1), C20(2), C20(3))
endfor
##################### end script ############################################
```

The four atoms were mapped onto the model with the FRAG...FEND command in SHELXL, e.g.

```
FRAG 17
C14 1       6.539   0.630   9.384
C15 1       7.650   0.766   8.614
C19 1       9.000   0.249   8.820
C20 1       9.298  -1.031   9.036
FEND
[...]
AFIX 170
C9     1    0.206012    0.214497    1.001840    11.00000    0.05895
C10    1    0.267544    0.166704    1.123857    11.00000    0.03763
C22    1    0.237086    0.125825    1.279921    11.00000    0.09427
C23    1    0 0 0    11.00000    0.07069
AFIX 0
```

Figure S4 gives an overview of the resulting dihedral angles of the vinyl groups after kinematical refinement of each starting model, either with (blue) or without (red) BUMP restraints. For the models of *R*/*S*′, the majority of refinements, especially those with BUMP restraints, there is a clear preference for the cis conformation of all vinyl groups. The refinements of *S*/*R*′ suggest that the parameter landscape has several local minima, and no clear preference is observed for the C′ vinyl group. In the case of *R*/*R*′, the refinements identify preferred dihedral angles for the vinyl groups of the B, C, and B′ pyrrole rings. Though the angles for the C′ vinyl group are more scattered, all of the refinements – including those that started with cis conformation – ended up with the C′ group in trans conformation.

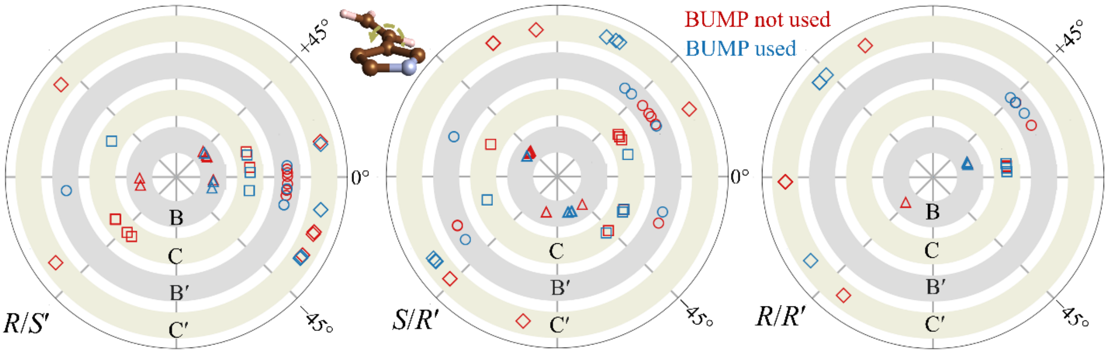

**Fig. S4**: Refined dihedral angles resulting from 8 starting models for each dimer. Vinyl groups were placed in such a way, that the initial dihedral angles adopted values between −180° and +180° in steps of 45°.

##### *Isomeric disorder in the model of R/S′ and R/R′*

As mentioned above, the disorder in the two methyl-vinyl groups of the lower hematin was modelled as an overlap of *R*′ and *S*′ of only the eight involved carbon atoms (and the related, constrained hydrogen positions). The rest of the molecule was modelled non-disordered. The two methyl-vinyl groups of the *S*′-isomer were placed into one part 1 with free variable 2, the two methyl-vinyl groups of the *R*′ isomer were placed into part 2 and constrained to the occupancy 1-fvar(2). The refined free variable fvar(2) thus corresponds to the occupancy of the *R*/*S*′-dimer. As occupancy and ADP values are strongly correlated, the ADPs of the atoms in the overlapping groups within one methyl-vinyl group were restrained, e.g. with

```
SIMU 0.01 0.02 C20_2^a C19_2^a C15_2^a C21'_2^b C15'_2^b
```

### 4. Dynamical Diffraction Analysis

#### Data selection and processing

Data sets with best data set statistics in terms of average reflection intensity <*I*/*σ*> and $CC_{1/2}$ were selected for the dynamical refinement. Of batch 1, the data sets with labels 03, 04, 07, 08, and 09 from day 1 and 01, 02, 07, and 08 from day 2 (https://doi.org/10.5281/zenodo.5039355) were transformed from MRC to TIFF format using the tool mrc2tif from IMOD [1]. Diffraction patterns were analysed with PETS2 [17]. Detector-specific noise parameters of the CMOS-type CETA-D detector were estimated to be 52 counts/electron for the gain (and cascade factor) and 11 counts for the variance of the readout noise. After determination of the unit cell parameters and initial indexing, the orientation of the frames was optimized, and distortions present in the diffraction patterns were corrected for. Unit cell constraints were not applied. An output file with integrated intensities, estimated uncertainties and relevant geometric parameters (orientation matrix, crystal orientation) was generated for each data set. For the calculation of dynamical integrated intensities, reflections were grouped into scaling and orientation batches (called virtual frames), each covering a goniometer rotation of 4° with a mutual overlap of 2°.

From batch 2, in total 13 data sets were selected and their diffraction patterns converted to TIFF format. The noise parameters were estimated to be 3 counts/electron and 1 count as variance of the readout noise. The remaining steps of the data reduction were performed as described for batch 1. Finally, 33 frames were merged to form one virtual frame, so that each covers a goniometer rotation of 3.63° with a mutual overlap of 1.21°.

#### Dynamical refinement with Jana2006

Nine output files based on data sets from batch 1 were imported to Jana2006 [18] and unit cell parameters were set to $a$ = 12.086 Å, $b$ = 14.6216 Å, $c$ = 7.9942 Å, $\alpha$ = 90.758°, $\beta$ = 97.093°, $\gamma$ = 97.060°. Electron scattering factors were taken from the International Tables for Crystallography [19]. The model with isomeric disorder from the kinematical refinement was used as a starting model. Soft restraints were used so that atomic distances and angles in the same local chemical environment are identical. Assuming that the hematin moieties behave like rigid bodies, displacement parameters were modelled with a set of translation, libration and screw (TLS) parameters. Translation and libration parameters of the *R* and *S*′ hematin were constrained to be equal, and screw parameters were set to 0. The methyl and vinyl groups at the α, β, α′ and β′ sites were not included in the TLS-description. Anisotropic displacement parameters of carbon atoms directly bonded to the B, B′, C and C′ pyrrole groups (C19, C21, C22, C24) were constrained to be equal. Isotropic displacement parameters of the remaining carbon atoms of the vinyl groups (C20, C23) were freely refined. There were in total 196 structural refinement parameters, including 12 TLS parameters and one occupancy parameter.

Additional refinement parameters specific for the dynamical refinement are one thickness parameter per data set (in total 9) and one scale parameter per virtual frame (in total 296). In the initial refinement cycle only frame scales and thickness parameters were refined. The subsequent refinement of all 501 parameters converged with $R_{obs}$ = 0.118 for 9656 observed reflections ($I/\sigma > 3$), $R_{all}$ = 0.247 and $wR_{all}$ = 0.146 for 27397 reflections. Details are given in Table S4 and the model was deposited at the Cambridge Crystallographic Data Centre CCDC (deposition number 2156990).

One occupancy parameter representing the fraction of the achiral *R*/*S*′ dimer was refined, which only affects the methyl and vinyl sites at the α′ and β′ sites (B′ and C′ pyrrole rings). The refined value converged to 0.528(9), suggesting that the fractions of achiral (*R*/*S*′) and chiral dimers (*R*/*R*′) are equal. For a subset of 6 data sets with the best refinement statistics (day 1: 07, 08, 09; day 2: 02, 07, 08), the sensitivity to the occupancy estimate was investigated by a set of refinements with fixed occupancy covering the range 0 to 1 (Fig. S5).

The reflection lists and further experimental parameters based on the 13 data sets of batch 2 were imported by Jana2006 and the same restraint and constraint scheme was used as described above. Refinements of single dimer models confirmed the observations from the kinematical refinements, i.e. suspicious displacement parameters in single dimer models indicate the presence of disorder. For the systematic investigation of the occupancy, displacement parameters of vinyl groups were constrained, otherwise only the displacement parameters of the overlapping atoms of the disordered methyl and vinyl groups were constrained. In agreement with the DFT results, models of *S*/*R*′ resulted in the worst fit ($wR_{all}$ = 0.120), and those of *R*/*R*′ ($wR_{all}$ = 0.104) and *R*/*S*′ ($wR_{all}$ = 0.105) were visibly better but still suffer from unrealistic displacement parameters. The best fit was obtained from a model representing a concentration of 56.3% of *R*/*S*′ dimers and 43.7% *R*/*R*′ dimers. The final model was deposited at the Cambridge Crystallographic Data Centre CCDC (deposition number 2240526).

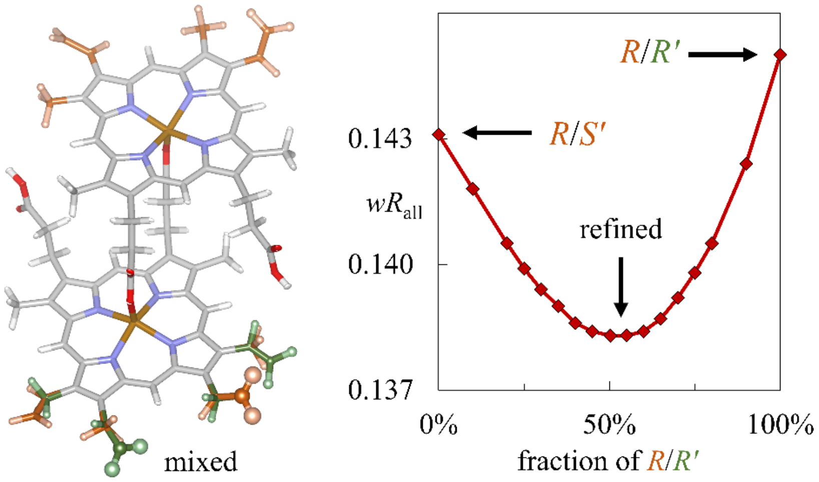

**Figure S5.** Left: Final model from the dynamical refinement against 9 data sets from batch 1 without constraints on the displacement parameters of the terminal carbon atoms of the vinyl groups. Right: Constraints on all displacement parameters were used for the systematic analysis of the dependence of the refinement *R*-factor on the (fixed) composition, again based on 9 data sets from batch 1. Qualitatively, these results are in very good agreement with results presented in Fig. 2.

| **Crystal data** | | | | | | |
|---|---|---|---|---|---|---|
| Chemical sum formula | $C_{34}H_{31}FeN_4O_4$ | | | | | |
| Molecule mass (Da) | 615.49 | | | | | |
| *Z* | 2 | | | | | |
| Crystal system, space group | triclinic, *P*1 | | | | | |
| a, b, c (Å) | 12.09(8), 14.62(9), 7.99(3) | | | | | |
| α, β, γ (deg.) | 90.8(3), 97.1(4), 97.1(4) | | | | | |
| *V* (Å³) | 1390(14) | | | | | |
| **Data collection** | | | | | | |
| Transmission electron microscope | Talos Arctica (Thermo Fisher Scientific) | | | Glacios (Thermo Fisher Scientific) | | |
| Acceleration voltage (kV), $\lambda$ (Å) | 200, 0.02508 | | | 200, 0.02508 | | |
| Sample temperature (K) | 80 | | | 80 | | |
| Data sets used | 9 | | | 13 | | |
| $h_{min}$, $h_{max}$ | −12, 12 | | | −14, 14 | | |
| $k_{min}$, $k_{max}$ | −15, 15 | | | −17, 17 | | |
| $l_{min}$, $l_{max}$ | −8, 8 | | | −9, 9 | | |
| **Data set statistics** | Overall | Low-resolution shell | High-resolution shell | Overall | Low-resolution shell | High-resolution shell |
| High-resolution limit (Å) | 0.93 | 2.46 | 0.93 | 0.83 | 2.17 | 0.83 |
| Low-resolution limit (Å) | 7.93 | 7.93 | 0.98 | 9.83 | 9.83 | 0.85 |
| Completeness (%) | 91.1 | 91.1 | 80.7 | 98.8 | 98.3 | 98.1 |
| Multiplicity | 4.2 | 4.0 | 3.6 | 4.9 | 4.3 | 3.3 |
| $<I/\sigma>$ | 4.7 | 19.3 | 1.5 | 2.7 | 13.6 | 0.39 |
| All reflections | 27397 | 1373 | 2746 | 48505 | 2430 | 2431 |
| Observed reflections | 9656 | 1051 | 436 | 10185 | 1648 | 22 |
| Total unique | | | | 9974 | 566 | 730 |
| **Dynamical refinement** | | | | | | |
| $RSg_{max}$, $DSg_{min}$ (Å$^{-1}$), $g_{max}$ (Å$^{-1}$) | 0.75, 0.0015, 1.1 | | | 0.7, 0.0025, 1.2 | | |
| No. refinement parameters | 501 | | | 666 | | |
| $R_{obs}$, $R_{all}$, $wR_{all}$ | 0.116, 0.245, 0.143 | | | 0.091, 0.198, 0.101 | | |
| $MR_{obs}$, $MR_{all}$, $MwR_{all}$ | 0.125, 0.215, 0.120 | | | 0.084, 0.148, 0.083 | | |

**Table S4.** Crystal structure determination applying dynamical diffraction theory.

#### Absolute structure

Although dynamical diffraction has been used for the determination of the absolute structure [20], in this case *R*/*R*′ and *S*/*S*′ hematin could not be distinguished at a satisfactory confidence level. 6 data sets of batch 1 (day 1: 07, 08, 09; day 2: 02, 07, 08) with the most promising refinement statistics were selected. The refinement of the model with *R*/*S*′ and *R*/*R*′ yields slightly better residuals numerically, but the result of this analysis remains ambiguous (Table S5). Likewise, the analysis of the 13 data sets of batch 2 did not allow to identify a clear preference for one enantiomorph or the other. There are several reasons why the assignment of the absolute structure may fail. First of all, the difference in intensities depends on the structural differences between the enantiomorphs, which in this case are limited to very few, partially occupied atoms. The simplistic modelling of the disorder leads to further deviations of the calculated intensities and worsens the fit. Furthermore, inelastic scattering is not taken into account and it is assumed that the crystal is perfect. Crystal defects and inelastic scattering are thus not only responsible for the relatively high *R*-factor, but in this case obstruct the determination of the absolute structure.

The models presented throughout the text assume that the crystal consists of *R*/*S*′ and *R*/*R*′ hematin dimers. This assignment of the absolute configuration is based on considerations regarding the crystal morphology and growth as described in the main text.

| Batch | Model | $R_{obs}$ | $R_{all}$ | $wR_{all}$ | Refined *R*/*S*′ fraction |
|---|---|---|---|---|---|
| 1 | *R*/*S*′ and *R*/*R*′ | 0.116 | 0.213 | 0.138 | 0.497(14) |
| | *R*/*S*′ and *S*/*S*′ | 0.117 | 0.212 | 0.139 | 0.541(13) |
| 2 | *R*/*S*′ and *R*/*R*′ | 0.092 | 0.198 | 0.101 | 0.553(6) |
| | *R*/*S*′ and *S*/*S*′ | 0.091 | 0.198 | 0.101 | 0.557(7) |

**Table S5.** Comparison of refinement residuals for the two enantiomorphs.

#### Additional views on the final crystal structure model

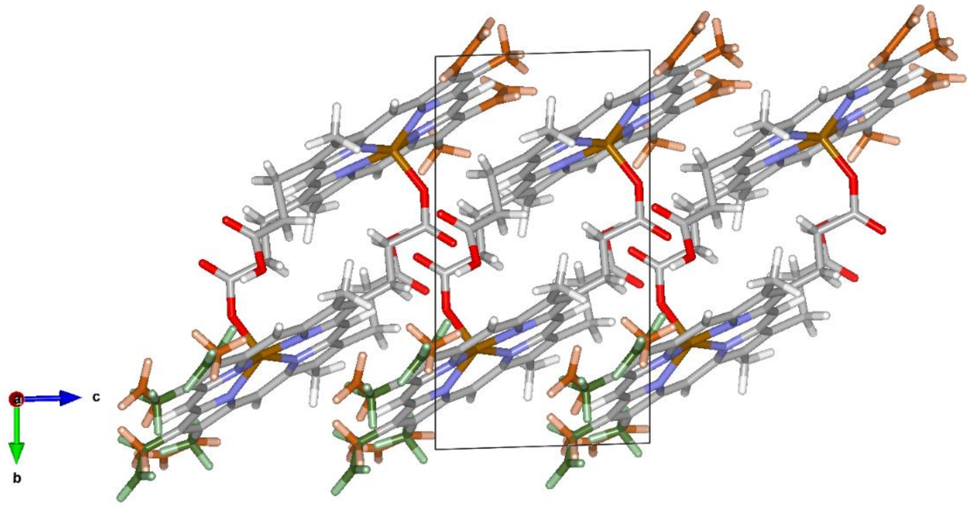

**Figure S6.** View along −**a**, i.e. [100] points towards the viewer, with disordered vinyl sites at the +**b** and −**c** side.

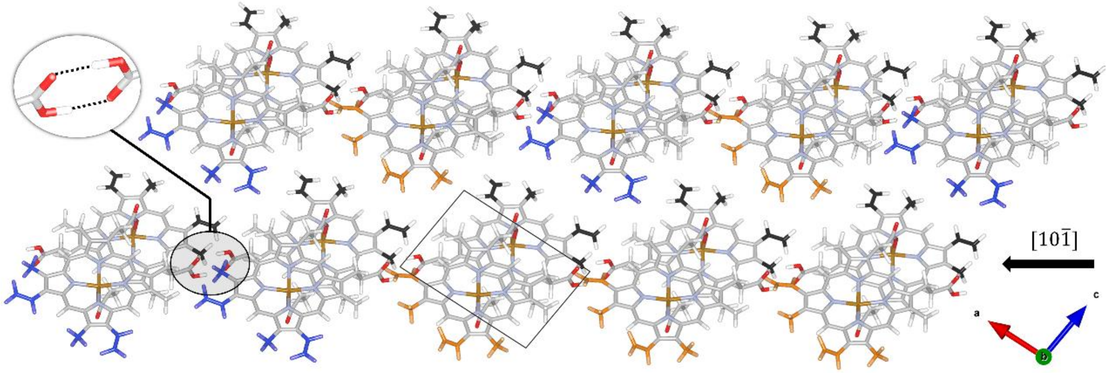

**Figure S7.** In the real structure, *R*/*S*′ and *R*/*R*′ dimers are distributed in a disordered way. Therefore, there are different local environment as neighboring dimers can be both of the same type or of different types. Independent of the disorder, hydrogen bonds form between the propionic acids leading to a chain of dimers along the $[10\bar{1}]$ direction. In this figure, methyl and vinyl groups of the B and C pyrrole rings are colored black. Methyl and vinyl groups of the B′ and C′ pyrrole rings are colored blue if their arrangement defines an *R*/*S*′ dimer, and colored orange if the arrangement defines an *R*/*R*′ dimer.

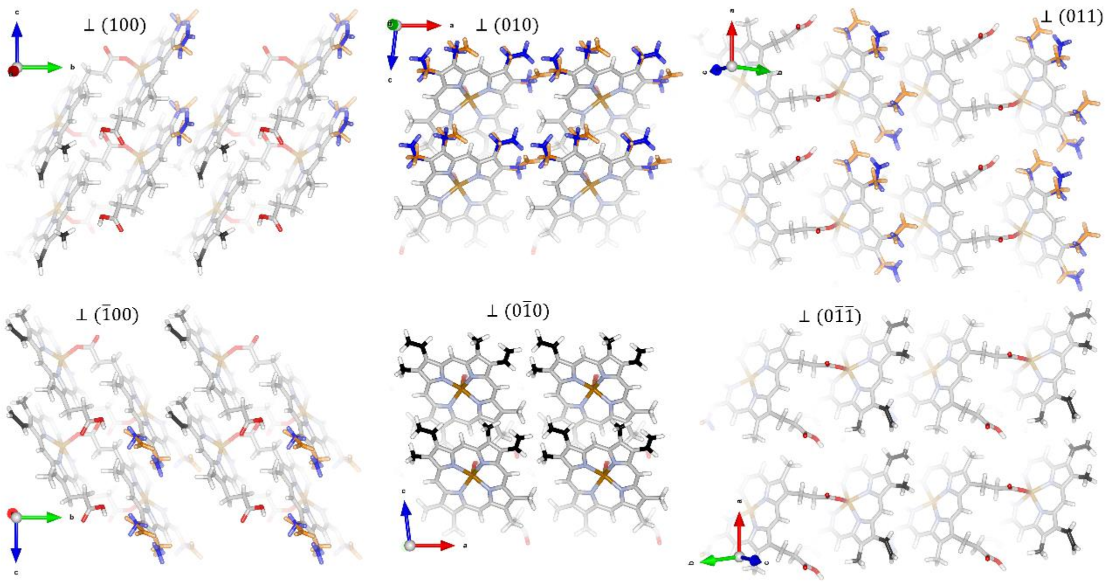

**Figure S8.** Crystal faces opposite to each other are not identical due to the chiral nature of hemozoin. Three pairs of opposing crystal faces with functional groups exposed at the surface are shown. Atoms fade out the farther away they are from the surface. Note that the morphology of the hemozoin crystals investigated in this study do not show clear signs of the $(011)$ and $(0\bar{1}\bar{1})$ faces.

#### 5. Density Functional Theory

##### Computational Details

DFT calculations were performed using the FHI-aims [21] code with *tier 2* numerical atomic-centered orbital (NAO) basis sets and tight numerical settings. The generalized gradient approximation (GGA) of Perdew, Burke, and Ernzerhof (PBE) [22] was combined with the many-body dispersion (MBD) [23,24] method. MBD has been shown to produce highly accurate lattice energies for molecular crystals [25–27]. In all DFT calculations, a 3x3x3 k grid was applied. The convergence criterion for geometry relaxations was a maximal force below $10^{-2}$ eV/Å. Initial structure models were based on results from the kinematical refinement in space group *P*1.

##### Analysis of vinyl conformation

The diffraction-based structure analysis revealed that, in addition to the large displacement parameters observed for two carbon atoms, one vinyl moiety of the chiral hematin dimer appeared to take a *trans* conformation while the other three were in *cis*. A search of the PDB found that in the majority of heme structures co-crystallized with heme binding proteins, both vinyl groups were in *cis* (https://dagewa.github.io/heme_explorer/). The *trans* vinyl was one of those with large displacement parameter for the terminal carbon in the diffraction-based structure analysis of single-dimer models. Therefore, we considered that crystal packing forces might influence its conformation. DFT calculations were conducted to test this hypothesis and to ascertain independently the positions of the vinyl groups on the chiral and *R*/*S*′ dimers.

For each dimer, a combinatorial scan was performed for all possible combinations of the vinyl groups in *cis* or *trans* conformation, a total of 16 configurations per dimer. For models comprising mixtures of *R*/*S*′ with *R*/*R*′ dimers, all 256 configurations were considered.

In the modelled crystal, the unit cell parameters were constrained to those determined experimentally by Straasø et al. at *T* = 80 K. Hence, only the internal atomic positions were relaxed. The relative energies of structures with different vinyl group configurations were then compared. Representative structures with their energies are provided in Figure S9. The energy of the centrosymmetric dimers *R*/*S*′ and *S*/*R*′ is lowered if all vinyl groups are in *cis* conformation without any *trans* vinyl group. For the *R*/*R*′ dimer the most stable configuration has a single *trans* vinyl group and 3 *cis* vinyl groups. In the mixed-dimer model, *R*/*S*′ adopts all *cis* vinyl groups while *R*/*R*′ has 1 *trans* and 3 *cis* vinyl groups (Fig. S9).

##### Relative stability of single-dimer and mixed-dimer models

The relative stability of single-dimer models (*R*/*S*′, *S*/*R*′, *R*/*R*′) and the mixed dimer model (1:1 mixture of *R*/*S*′ and *R*/*R*′) was evaluated with a full unit cell relaxation (Table S6, Figure 3). The stability of a 1:1 mixture of *R*/*S*′ and *R*/*R*′ (or *S*/*S*′) agrees very well with the formation of biogenic hemozoin 1:1 mixture of *R*/*S*′ and *R*/*R*′. Similar to the analysis of the vinyl conformation with fixed unit cell parameters, the calculations with relaxed unit cell parameters confirm the identified vinyl orientations, especially the single *trans* vinyl group of *R*/*R*′ dimers (Figure 3, Table S6). The DFT calculations thus further support the findings of the electron diffraction analysis which identified the same vinyl group to be in a *trans* conformation.

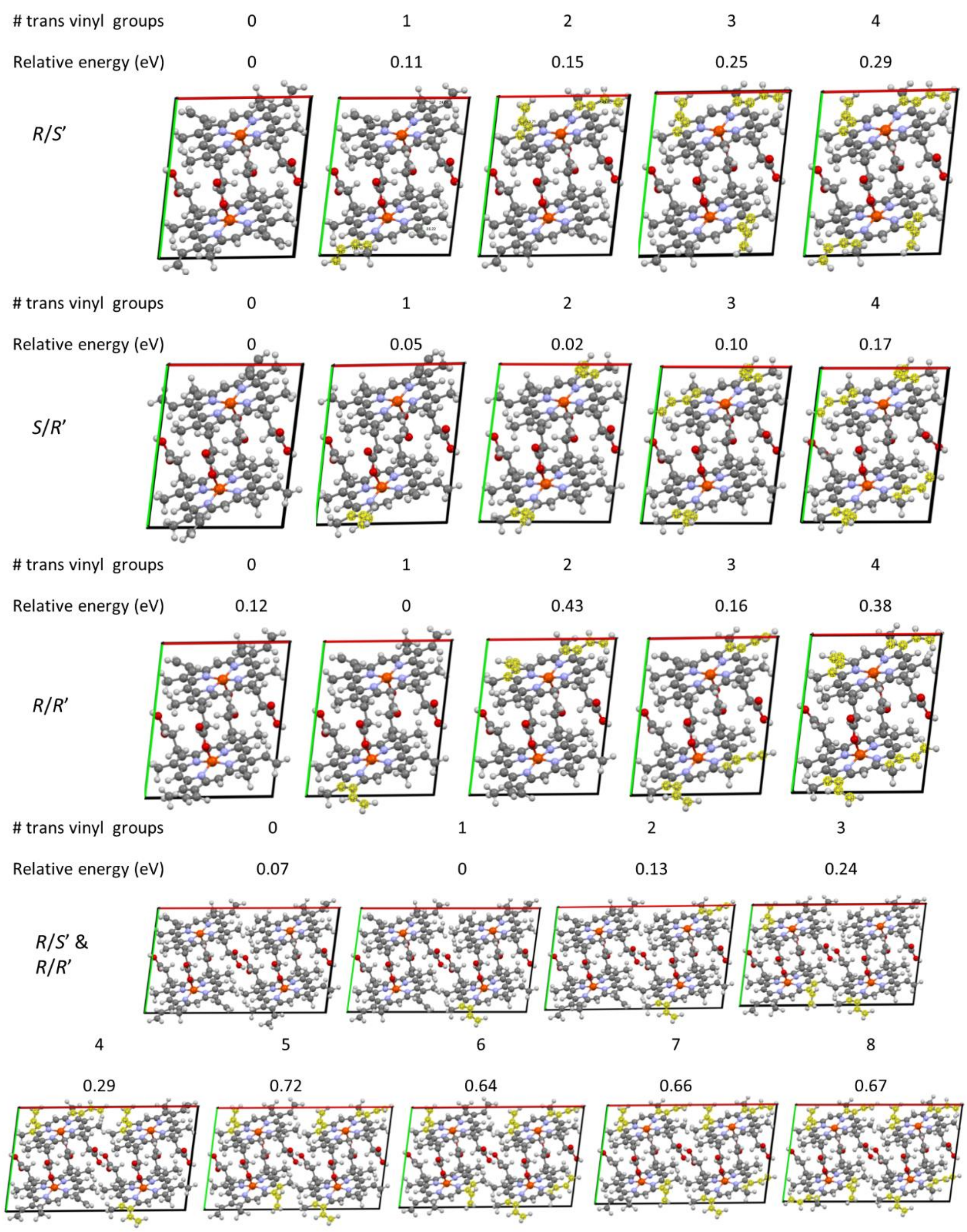

**Figure S9.** Representative structures from combinatorial structure search of *R*/*S*′, *S*/*R*′, *R*/*R*′ and the mixture of *R*/*S*′ with *R*/*R*′. The dependence of the energy on the number of trans vinyl groups is shown. Results are obtained with PBE + MBD with tight tier2 numerical atomic-centered orbital (NAO) basis set. The most stable configuration is referenced to zero.

#### Comparison of unit cell parameters and selected distances

Relaxed unit cell parameters of *R*/*S*′ and *R*/*R*′ in this work are similar to those reported by Straasø et al. at $T$ = 80 K, while the relaxed unit cell parameters of *S*/*R*′ deviate visibly, especially $b$ and $\gamma$. Structural parameters from a previous DFT study are not directly comparable because different unit cell parameters were used without relaxing them and a different dispersion-correction method was used [28]. Despite those differences, the Fe-O and Fe-Fe distances in this work are both closer to values determined experimentally by electron diffraction (Table S6).

| | DFT | | | | | ED (batch 1) | | ED (batch 2) | |
|---|---|---|---|---|---|---|---|---|---|
| **Label** | ***R/S′*** | ***S/R′*** | ***R/R′*** | ***R/S′* + *R/R′*** | | ***R/S′* + *R/R′*** | | ***R/S′* + *R/R′*** | |
| relative E (eV) | 0.00 | 0.27 | 0.13 | 0.06 | | *not applicable* | | *not applicable* | |
| **unit cell parameters** | | | | | | | | | |
| *a* (Å) | 12.08 | 12.12 | 12.11 | 12.15 x 2 | | 12.086 (4) | | | |
| *b* (Å) | 14.29 | 15.11 | 14.40 | 14.20 | | 14.6216 (5) | | | |
| *c* (Å) | 7.95 | 7.95 | 7.97 | 7.97 | | 7.9942 (4) | | | |
| *α* (deg.) | 91.3 | 91.8 | 91.1 | 91.6 | | 90.758 (4) | | | |
| *β* (deg.) | 96.4 | 98.2 | 98.0 | 96.6 | | 97.093 (4) | | | |
| *γ* (deg.) | 97.1 | 100.6 | 96.7 | 96.5 | | 97.060 (5) | | | |
| *V* (Å³) | 1353.2 | 1415.3 | 1365.5 | 1356.3 x 2 | | 1390.1 | | | |
| **selected distances (Å)** | | | | | | | | | |
| *d*(Fe-Fe) | 9.02 | 9.05 | 9.02 | 9.00, 8.97 | | 9.05 | | 9.03 | |
| *d*(Fe-O) | 1.85 | 1.85 | 1.84 | 1.84, 1.84 | | 1.90, 1.92 | | 1.87, 1.90 | |
| **vinyl dihedral angles (deg.)** | | | | | | | | | |
| *φ*(B) | 28.45 | -24.53 | 29.44 | 25.87 | 27.92 | 32.3 | | 31.4 | |
| *φ*(C) | 23.90 | -21.34 | 23.31 | 25.39 | 21.48 | 15.4 | | 16.9 | |
| *φ*(B′) | -28.45 | 24.53 | 24.71 | -41.08 | 15.72 | 3.6 | 22.7 | 5.3 | 29.2 |
| *φ*(C′) | -23.90 | 21.33 | -143.62 | -5.77 | -144.64 | 16.5 | -152.6 | 13.7 | -139.3 |

**Table. S6.** Relative energy per dimer (most stable structure is referenced to 0), unit cell parameters, selected distances, and vinyl torsion angles of geometrically optimized structure models, obtained with PBE + MBD with tight tier2 numerical atomic-centered orbital (NAO) basis set. Relevant parameters from the dynamical refinement against electron diffraction data (label ED) are also included. Note that the unit cell parameters used in the refinement were taken from Straasø et al. (2014). Labels of the dihedral angles *φ* refer to the pyrrole ring to which the vinyl group is bonded. The calculation of the dihedral angle is based on four C atoms, of which two belong to the vinyl group and the other two are the carbon atoms of the pyrrole ring between the vinyl group and the closest methyl group. For the mixed model, the dihedral angles of the *R*/*S*′ dimer appear in the left column, while those of the *R*/*R*′ dimer are in the right column. The *trans* dimer is highlighted with yellow background.

#### 6. High-resolution TEM and Focal Series Reconstruction

In contrast to kinematic diffraction analysis, chiral handedness can in principle be resolved by real space imaging at atomic resolution. HRTEM images were recorded on a cryo-sample of dispersed hemozoin crystals in a Titan Krios G3 microscope (Thermo Fisher Scientific Electron Microscopy Solutions, Hillsboro, USA) at an acceleration voltage of 300 kV. To avoid image interpretation artefacts produces by lens aberrations during the imaging process we employed focal series reconstruction (FSR) (Thust, 2016) to obtain a representation of the exit-plane wave-function (EPWF) that emerges as a consequence of the interaction between the incident electron wave and the hemozoin crystal. The phase of the EPWF is a direct representation of the projected electrostatic potential of the crystal in the case of a sufficiently thin object, thus providing unfalsified structure representation. In this case we took advantage of low-dose FSR [29], a technique employing fast focal series on a low dose counting detector. The objective lens of the Krios TEM was ramped in discrete focus steps of 5 nm while a fast series of hundreds of frames were recorded on a Gatan K3 direct electron detector (Gatan Inc., Pleasanton, USA). Counting mode was used with a frame rate of 13 ms at the highest duty cycle. The fluence was 7 $e^{-}$/pixel/s and the series lasted a total of 4 seconds. The focal range for series acquisitions was between Lichte focus and Scherzer focus [30], i.e., between minimum delocalization and maximum point resolution focus. Low-dose conditions were used for object selection at low magnification and focus adjustment on an area adjacent to the crystal. A fast Fourier transform was computed immediately, from which the lattice parameters and a rough estimate of the orientation could be determined.

From many dozens of crystals, a total of 26 recordings were taken. Of those, only a single one provided a high resolution view in the desired direction along the *a* axis where the atomic columns of the methyl-vinyl groups are visible. Post-processing, this crystal was further analyzed by diffraction tomography in a Talos Arctica microscope at 200 kV. Diffraction patterns were recorded on a Gatan OneView IS camera and analyzed using DIALS by the same pipeline described above in order to establish the absolute lattice orientation, which remains ambiguous in a projection image. The axis orientations shown reflect this analysis.

A custom-written Python program [29] was used for the post-acquisition EPWF reconstruction, using raw data from selected subframes of the entire field of view of the K3 camera as input. The custom code implements a linear reconstruction scheme, following the detailed description by Thust and Kübel [31,32]. Nonlinear optimization schemes were not applied to the sparse low-dose data because of stability constraints on the optimization algorithm. The EPWF was refocused to the object plane through a propagator.

Results appear in Figure S10. The experimental data shown in Figure S10a-c was obtained from a full series of images over 4 s with a cumulated fluence 160 $e^{-}/Å^{2}$. A thin area at the edge of the crystal was selected for reconstruction, and the phase of the exit-plane wave was reconstructed to a resolution of 5 $nm^{-1}$. The phase of the refocused EPWF was averaged over a 5x5 lattice to reduce noise in the individual centers. It was immediately apparent that there are two different types of channels, one “filled” and one relatively “empty” (labelled with 1 and 2 in Fig. S10c), alternating in position. By overlaying the atomic models, it was clear that the filled holes contain the propionic acid bridges, and that the apparently empty channels actually represent the area of interest containing juxtaposed methyl-vinyl groups. One of these should be the “upper” (unprimed) monomer and the other the “lower” (primed) monomer of the neighboring dimer. Moreover, the density within the channel appears not to be symmetric, with a spike or shpitz often emerging from below in the image.

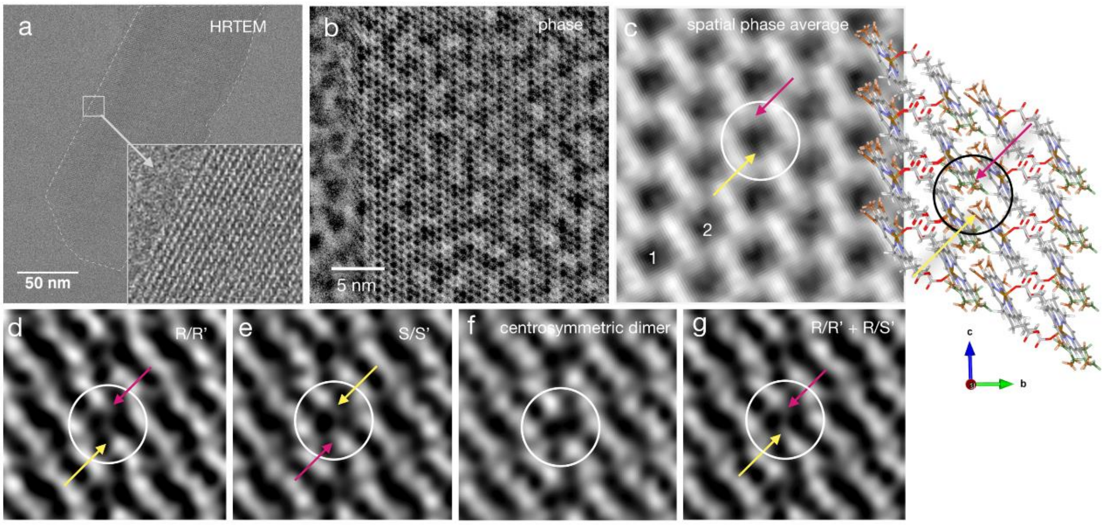

**Figure S10.** Low-dose focal series reconstruction of a hemozoin crystal. (a) Single image of a hemozoin crystal, taken from a through-focus series. Titan Krios, K3 counting mode, total fluence: 160 $e^-/Å^2$ for 300 images. (b) Phase of the exit-plane wave function, reconstructed from 240 images. Reconstruction limit is 5 $nm^{-1}$. (c) Magnified view of the reconstructed phase, running average over 5x5 repeat cells, Fourier-filtered at 4 $nm^{-1}$. Two distinct channels align vertically, marked by 1 and 2. Propionic acid bridges are located in channel 2. The orientation of the high-density channels with the propionic acid bridges with respect to the crystal edge is in agreement with the −**a** viewing direction of the crystal. In channel 1 an asymmetric contrast is observed, with a shpitz feature on one side (yellow arrow) and a fist feature on the opposite side (magenta arrow). (d) - (g) Calculated phase images, obtained by forward simulation of focal series images from four structure models, and subsequent focal series reconstruction with reconstruction limit of 5 $nm^{-1}$ and subsequent spatial period averaging over 3x3 repeat cells. The centrosymmetric structure fails to reproduce the asymmetric phase distribution (shpitz and fist) observed in the channels of type 1. The refined structure models with displacement order reproduce the fist feature on the side of the higher disorder. The refined structure with *R*/*R*′ and *S*/*S*′ configuration reproduces the experimental asymmetry in the phase image well. A structure model representing the lattice containing the mixture of *R*/*R*′ and *R*/*S*′ dimers is displayed to the right. The disorder appears on the first side (magenta arrow in the circular marker).

In order to interpret this asymmetry, we used the atomic models generated from the diffraction data (Tab. S4) to simulate forward the phase images that would ideally be obtained using the Krios microscope. The first step in such a forward simulation was the creation of an atomistic super-cell of a slab-like hemozoin crystal from unit cell data by periodic repetition. A slab of a fully atomistic model of amorphous carbon was added, representing the support film of the TEM grid. In detail, the thickness of the hemozoin crystal and the amorphous carbon support were refined to match experimental images resulting in a final thickness of 20 nm for the hemozoin and 7 nm for the support. In the second step of the simulation a multislice calculation was used to generate the ideal EPWF and a focal series of images. These multislice calculations were carried out with the command line tools for TEM simulation included in the Dr. Probe simulation package [33]. Simulated focal series images were then transformed to counting mode images that approximate a Poisson-distribution of the number of electron counted in each pixel of the image for an accumulated number of frames, with an expectation value according to the electron flux in experiment. The simulated low-dose counting mode images were then used in the third step of the forward simulation as input for the LD-FSR reconstruction algorithm, with identical reconstruction parameters as for the experimental images.

Results of this forward simulation are shown in Figure S10d-g for four cases, a pure chiral *R*/*R*′ (Fig. S10d) and *S*/*S*′ model (Fig. S10e), a centro-symmetric dimer model of *R*/*S*′ (Fig. S10f) and the refined model that contains a mixture of chiral *R*/*R*′ dimers and *R*/*S*′ dimers (Fig. S10g). The simulated images of the chiral dimer arrays and the mixture of *R*/*R*′ and *R*/*S*′ dimers revealed an asymmetric protrusion of density into the channel (Figure S10d, e and g). For *S*/*S*′ the spike intruded from the top, and for *R*/*R*′ from the bottom in the orientation shown. The channel densities appeared symmetric for the centro-symmetric models (Figure S10f), as expected. The disorder appears on the fist side, where the larger effective displacement factors are associated with a delocalized contrast. Therefore, the results of this real space analysis are consistent with a contribution of *R*/*R*′ rather than *S*/*S*′. This assignment is in agreement with and further supports the absolute structure determination based on the crystal morphology.

#### 7. Model for oriented dimer formation by Heme Detoxification Protein

The HDP sequence was submitted to structure prediction servers including Phyre2 [34], I-Tasser [35], and the recent RoseTTAFold [36]. Additionally, a model became available at the AlphaFold Protein Structural Database [37,38], for UniProt ID A0A144A6G1. There was a broad consensus in the architectural features (Fig. S11). The N terminus up to amino acid 50 is almost entirely disordered, followed by a compact α-helical domain until aa125. The C terminal half includes prominent unfolded loops, with a single β-sheet appearing from aa188 near the end. The servers provided essentially identical predictions for the secondary structures, with significant variability in the orientation of the loops. After excluding the first 50 amino acids from the AlphaFold result, only 11 residues fall below a confident model prediction of pLDDT > 70. These residues form the loop from Ser137 to Lys147. Similarly, after excluding the first 50 amino acids from the RoseTTAFold result, this loop is the only region with an estimated Cα RMS error of greater than 3.0 Å. When that loop is also excluded, the mean estimated RMS error is 1.1 Å. We used these models to locate the two heme binding sites that had been identified previously by biochemical methods [39,40]. Most strikingly, the site spanning His122 and His197 appears in a pocket between the strongly and weakly folded domains. Presuming that the hematin Fe binds to His122, as found earlier by spectroscopy [41], and one of the propionic acid tails binds to His197, the vinyl/methyl pairs are oriented deeply in the protein structure where chiral discrimination is likely to take place. The other heme binding site consisting of His172 and His175 appears at the tip of an unfolded loop. These histidines were not found to bind the Fe.

The predicted structure suggests a model of dimer formation that limits the possible number of isomers formed. Specifically, the compact N terminal domain serves as a compact core, or “pedestal”, and the C terminal loops act as a loose cap. One hematin monomer anchors to the pedestal via the Fe-His122 interaction, with a planar orientation provided by His197. Steric hindrance between the vinyl-methyl end and the α-helices restricts the orientation around the long axis of the heme so that the histidine binds from the same side. Indeed it is a common feature of heme binding proteins that the binding seen in co-crystallization is oriented uniquely one way or the other [42]. The other heme binding site shows no suggestion of orientational preference, nor of a fixed spatial position. If the latter gets close to the orientation-sensitive binding site, this would allow the two hematin monomers to dimerize. Alternatively, reversible binding to one or the other of His172 and His175 might serve simply to raise the local concentration. With the orientation of one hematin already fixed by the protein pedestal, only one chiral isomer could be produced. Centro-symmetric dimers would be produced as well. In principle both *R*/*S*′ and *S*/*R*′ isoforms could be generated (Fig. S12). Note however, that the diffraction data analysis as well as the DFT results suggest that the *S*/*R*′ dimer would crystallize with unit cell parameters different to those experimentally observed (Fig. 2 and 3). Whichever side the fixed monomer exposes, there could be a steric hindrance to approach the other monomer from the left or the right. This would imply a kinetic preference for the formation of one or the other centrosymmetric dimer. Details of the HDP model are necessarily speculative since they depend on homology modelling and structure prediction, but the proposition is topologically robust: if the binding of one monomer is uniquely oriented, then only one chiral dimer is permitted (Fig. S12).

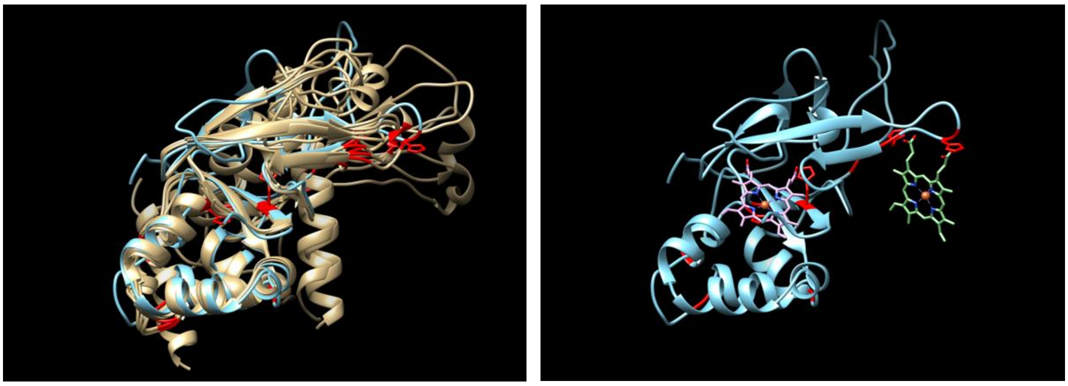

**Figure S11.** Structural modeling of the homologous protein Fascilin-1 of Drosophila (Uniprot code P10674) reveals a two-domain protein with a compact alpha-helical fold at the N terminus and a largely unfolded domain at the C terminus. The very end of the C terminus comes back to contact with the compact core. Left: Five models produced by RosettaFold are shown in beige, with one model produced by the Phyre2 server in teal. The overlap is very high in the N terminal region, and weaker in the C terminal region except for the two β-sheets. Histidines are indicated in red, and the key histidines 122, 197, 172, and 175 are shown as stick models. Right: The heme binding sites are shown on the single Phyre2 model (for clarity). One heme fits in a pocket formed between the compact and loosely folded domains, with His122 in proximity to the Fe as found previously by spectroscopy [41]. The fit is manual and only suggestive in details, but so long as the heme fits in only one orientation (purple), only one chiral dimer can be formed by the addition of the other monomer (green) from above. The heme binding site at the end of the unfolded loop shows no indication of an orientational preference.

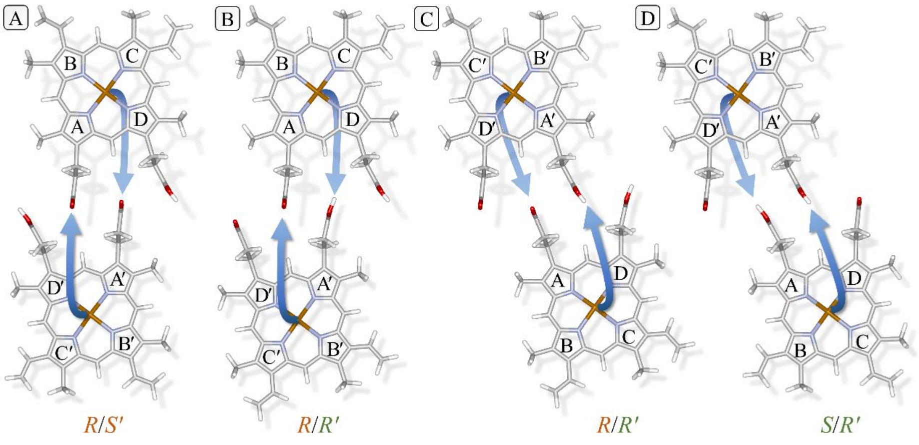

**Figure S12.** If one monomer has a fixed orientation (top row, above the plane), the choice of the orientation (two possibilities) and propionic acid bonding to Fe (two possibilities) of the other monomer (bottom row, below the plane) does not allow the formation of *S*/*S*′. Dimerization with two different orientations of the bottom monomer with its Fe bonding to the left propionic acid leads to the formation of (A) *R*/*S*′ and (B) *R*/*R*′. Bonding of Fe to the other propionic acid leads to the formation of (C) *R*/*R*′ and (D) *S*/*R*′. Note that in (C) and (D) the formed dimers must be rotated by 180° around the horizontal axis to get to the orientation as used in Figs. 1 and S3, leading to the shown assignment of pyrrole labels.
